# Beyond inappropriate fire regimes: a synthesis of fire-driven declines of threatened mammals in Australia

**DOI:** 10.1101/2022.03.15.483873

**Authors:** Julianna L. Santos, Bronwyn A. Hradsky, David A. Keith, Kevin Rowe, Katharine L. Senior, Holly Sitters, Luke T. Kelly

**Affiliations:** School of Ecosystem and Forest Sciences, The University of Melbourne; Centre for Ecosystem Science, School of Biological, Earth and Environmental Sciences, The University of New South Wales, Sydney, Australia; New South Wales Department of Planning, Infrastructure and Environment, Parramatta, Australia; Sciences Department, Museums Victoria, Melbourne, Australia; School of BioSciences, The University of Melbourne

**Keywords:** biodiversity, demographic processes, dispersal, extinction, fire frequency, movement, reproduction, survival, wildfire

## Abstract

Fire can promote biodiversity but changing patterns of fire threaten species worldwide. While scientific literature often describes ‘inappropriate fire regimes’ as a significant threat to biodiversity, less attention has been paid to the characteristics that make a fire regime inappropriate. We go beyond this generic description and synthesize how inappropriate fire regimes contribute to declines of animal populations, using threatened mammals as a case study. We developed a demographic framework for classifying mechanisms by which fire regimes cause population decline, and applied the framework in a systematic review to identify fire characteristics and interacting threats associated with population declines in Australian threatened land mammals (n=99). Inappropriate fire regimes threaten 88% of Australian threatened land mammals. Our review indicates that intense, large, and frequent fires are the primary cause of fire-related population declines, particularly through their influence on survival rates. However, several species are threatened by a lack of fire and there is considerable uncertainty in the evidence base for fire-related declines. Climate change and predation are documented or predicted to interact with fire to exacerbate mammalian declines. This demographic framework will help target conservation actions globally and would be enhanced by empirical studies of animal survival, dispersal, and reproduction.

## 1 INTRODUCTION

Fire is an important ecological process that can promote biodiversity (Jones & Tingley, 2021). Yet human actions are transforming fire activity and at least 4,400 species worldwide face threats associated with changing patterns of fire (Kelly et al., 2020). This includes 16% of all mammalian species classified as threatened with extinction by the *International Union for Conservation of Nature* (IUCN) (Kelly et al., 2020). While numerous research papers and policy documents describe ‘inappropriate fire regimes’ as a major threat to biodiversity, the specific characteristics that make a fire regime inappropriate receive less attention. Understanding the mechanisms through which inappropriate fire regimes cause population declines is critical for addressing biodiversity loss (McLauchlan et al., 2020) and is likely to create opportunities for more effective conservation actions (Nicol et al., 2019).

Four main factors make identifying the characteristics of inappropriate fire regimes difficult. First, fire regimes involve multiple components, including fire frequency, intensity and season (Gill, 1975), and their spatial dimensions. Second, population declines associated with fire regimes can be caused by a range of mechanisms that directly and indirectly influence animal survival, colonization, and reproduction (Whelan et al., 2002). Third, fire regimes and their impacts on populations may be a consequence of interactions between fire and other processes such as climate change (Hale et al., 2016), grazing (Probert et al., 2019), habitat fragmentation (Driscoll et al., 2021) and predation (Hradsky, 2020). Finally, the diverse life-history characteristics and habitat requirements among biotic communities lead to a variety of responses to fire regimes within (Senior et al., 2021) and between (Jones & Tingley, 2021) ecosystems.

Demographic approaches offer a way forward. Considering the effects of fire on key processes that shape populations – survival, reproduction and movement (Begon & Townsend, 2020) – provides a way to identify the mechanisms that underlie fire-driven population declines and forecast population changes. For example, Miller et al. (2019) developed a demographic approach to synthesize fire seasonality effects on plant populations and explore traits that make species vulnerable to decline. To date, demographic approaches for application in fire-prone ecosystems have primarily been developed to explore the ecology of plants (Keith, 1996; Miller et al., 2019). While important steps have been taken to develop conceptual demographic frameworks for animals (Whelan et al., 2002), they have not been applied systematically to assess threats related to inappropriate fire regimes at a continental scale.

Our overarching aim is to go beyond inappropriate fire regimes as a generic descriptor of threatening processes by applying a demographic approach to synthesize *how* fire regimes influence populations of threatened animals. We do this using a case study of Australian land mammals, a distinctive and mostly endemic group of species that continues to suffer high rates of population decline (Geyle et al., 2018; Woinarski et al., 2015). More than 300 terrestrial mammalian species are native to Australia, but their recent and ongoing declines are exceptionally high with over 10% of species extinct since European colonization (Woinarski et al., 2015). A further 99 taxa (species, subspecies, or populations) are listed as threatened with extinction under the Australian Government’s key environmental legislation, the *Environment Protection and Biodiversity Conservation Act 1999* (*EPBC Act*). Disentangling the causes of Australian mammalian declines has proved difficult, but a consensus is emerging that multiple drivers are in play, including modification of fire regimes, predation by introduced mammals, and habitat loss and fragmentation (Doherty et al., 2015; Fisher et al., 2014; Johnson, 2006; McKenzie et al., 2007).

We asked three main questions: 1) What are the underlying mechanisms by which inappropriate fire regimes cause population decline?, 2) Which characteristics of fire regimes, on their own or via interactions with other processes, are associated with population decline of Australia’s threatened mammals?, and 3) Are threats posed by inappropriate fire regimes associated with particular mammalian taxonomic groups or ecosystems?

We expect this demographic approach will be useful for assessments of other fauna in fire-prone ecosystems across the globe, as well for developing policies and actions to conserve Australia’s distinctive mammalian fauna.

## 2 DEMOGRAPHIC APPROACH AND SYSTEMATIC REVIEW

In summary, we identified three groups of demographic processes that are fundamental to changes in population size – survival (deaths), reproduction (births) and movement into and out of a population (immigration and emigration) (Begon & Townsend, 2020) – and seven mechanisms by which fire may impact these processes (Figure 1). We then systematically reviewed conservation assessments (two comprehensive sources) and primary literature on Australian terrestrial mammals listed as threatened under the *EPBC Act*, and applied the demographic framework to identify which characteristics of fire regimes, and interacting abiotic and biotic processes, are associated with the seven fire-driven mechanisms of population decline. The demographic framework is informed by Keith (1996), who identified mechanisms of fire-related population declines of plants, and Whelan et al. (2002), who emphasized critical life-cycle processes that relate to fire-driven population changes.

**FIGURE 1.**
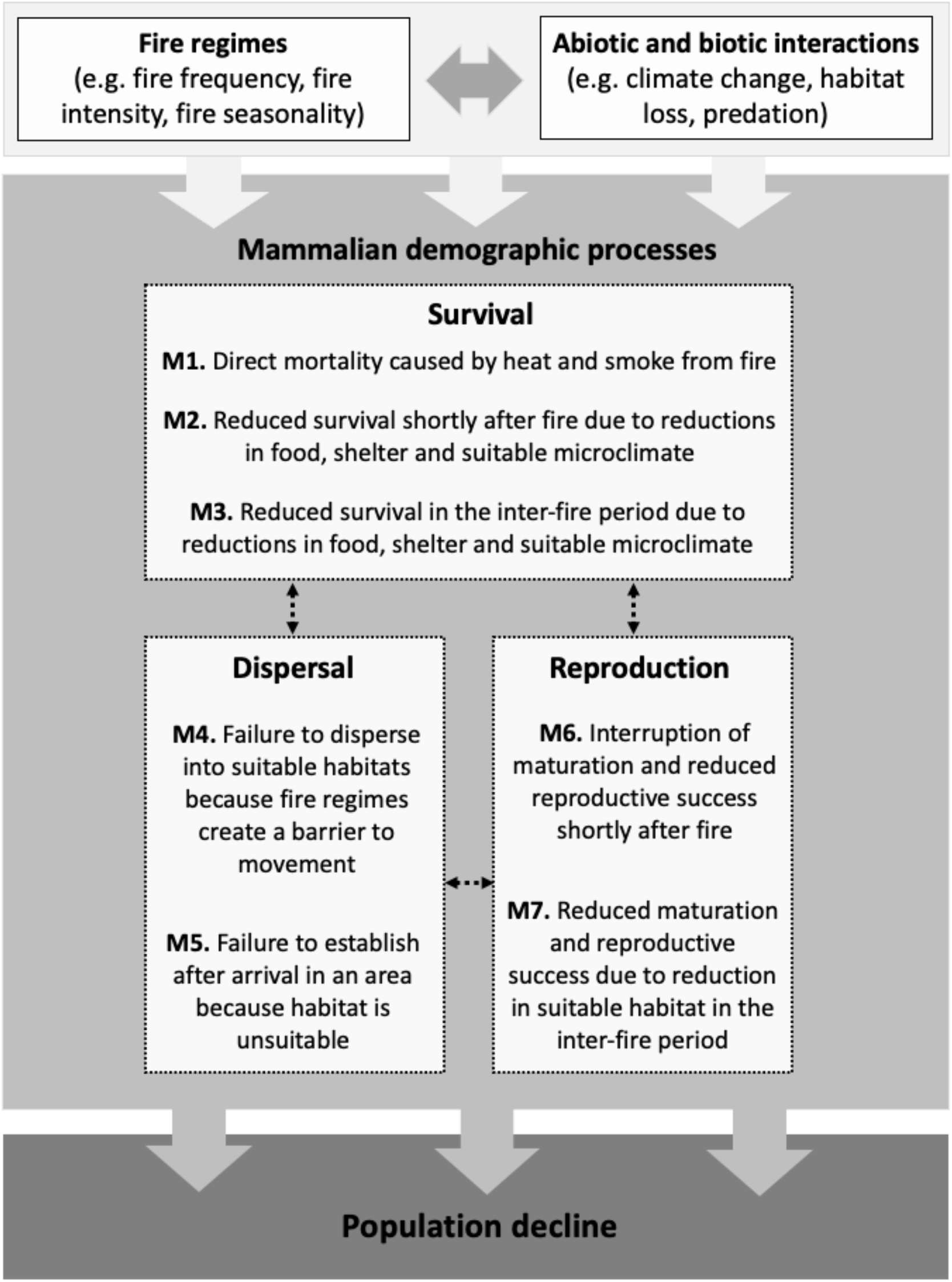
A demographic framework for assessing fire-driven mechanisms of population decline and extinction. Fire regimes and their interactions with other abiotic and biotic processes can negatively impact three demographic processes – survival, movement, and reproduction – via seven primary mechanisms (M1 – M7), leading to mammalian population decline and extinction. Arrows between the dashed boxes represent relationships between demographic processes and, in turn, these processes influence population change. A population may decline if fire reduces one or more of these demographic processes, and extinction occurs when the number of individuals declines to zero. The timing of each mechanism in relation to fire events or recurrent fire varies among species, and populations may decline because of one mechanism or a combination of mechanisms depending on a species’ life-history and habitat preferences.

We define inappropriate fire regimes as patterns of fire that may plausibly cause one or more taxa to decline. In many cases, we expect that inappropriate fire regimes result from modified patterns of fire. But it is possible that historical fire regimes applied under new environmental conditions could also cause mammalian populations to decline.

### 2.1 Description of fire-driven mechanisms of mammalian decline and extinction

#### Survival: Mechanisms 1, 2 and 3

Fire can cause population decline by reducing the survival of individual mammals in three main ways. First, fire can directly kill animals through exposure to high temperatures or smoke (Mechanism 1) (e.g. Koprowski et al., 2006; Silveira, 1999). Second, animals may survive a fire event but experience reduced rates of survival in the days, weeks and months after fire due to the depletion of food or shelter resources in recently burned areas (Mechanism 2) (Morris et al., 2011). Resource depletion can result from a single fire event or the cumulative impacts of multiple fires (Pardon et al., 2003), and operate at local (e.g. 1-10s ha), landscape (e.g. 1,000s-10,000s ha) and regional (e.g. 100,000s ha) scales. Third, fire-related reductions in survival can occur over years and decades if there is a decline in the quality of habitat and crucial resources following long intervals without fire (M3) (Arthur et al., 2012; Sherman & Runge, 2002). For example, some species benefit from early or mid-successional habitats that may become unsuitable after long periods without fire (Hayward et al., 2005). This could be a result of the senescence of food plants or adverse changes in vegetation structure and shelter availability.

The spatial dimensions of fire regimes can mediate Mechanisms 1, 2 and 3, through the effects of internal fire refuges (Shaw et al., 2001; Robinson et al., 2013) and habitat complementation (Kelly et al., 2017). Interactions with other processes, including predation, can also influence mammalian survival after fire (Hradsky, 2020). Here we group mechanisms of decline based primarily on demographic processes, and later link fire-regime characteristics and interacting threats to these mechanisms through systematic review.

#### Movement: Mechanisms 4 and 5

Movement within and between habitat patches of differing fire histories is a key determinant of animal distribution and abundance in fire-prone ecosystems (Nimmo et al., 2019). Colonization is a particularly important process involving the movement of animals, and includes dispersal to and establishment in new locations (or in locations where animals were previously present, called recolonization) (Hanski, 1999). We identify two mechanisms of decline relating to colonization and recolonization: failure to disperse into suitable habitats because fire regimes create a barrier to movement (Mechanism 4) and failure to establish after arrival in an area because habitat is unsuitable (Mechanism 5). Failure to disperse into suitable habitats (Mechanism 4) may be mediated by fire regimes that change the connectivity of habitats (Banks et al., 2013). A barrier to movement could be plausibly created by recently burned vegetation (Banks et al., 2015) or long unburned vegetation (Gavin et al., 1999; Pereoglou et al., 2013), depending on the species’ life-history and habitat requirements. Failure to disperse can result in population decline due to reductions in population size or gene flow (Sherman & Runge, 2002).

If dispersing animals arrive in a given area, colonization may still fail if they are unable to establish (Mechanism 5). Similar to barriers to dispersal, establishment limitation due to lack of resources could plausibly occur in recently burned areas or long unburned areas, depending on the species’ habitat requirements (Woinarski et al., 2005).

#### Reproduction: Mechanisms 6 and 7

Successful reproduction by surviving individuals or dispersing colonists is essential for the recovery of populations after fire events and the persistence of populations under recurrent fire (Whelan et al., 2002). In the short-term, fire can directly reduce reproductive success by interrupting the recruitment of mature individuals into the breeding population, delaying breeding by already mature individuals or reducing the availability of food or nesting resources needed for successful reproduction (Mechanism 6) (Griffiths & Brook, 2015). This is more likely if fire occurs during the breeding season (Morris et al., 2011) or when individuals are at vulnerable life stages (e.g. juvenile) (Laurenson, 1994).

Maturation and reproductive success can also decrease if habitat suitability declines in the inter-fire period, i.e., when there are long intervals without fire (Mechanism 7). Reduced maturation and reproductive success may occur if the extent or carrying capacity of suitable habitat decreases in response to long-term fire exclusion (Sherman & Runge, 2002).

### 2.2 Threatened species data and assessment of fire-related threats

We conducted a systematic review of the 99 Australian terrestrial mammalian taxa (including species, subspecies, and distinct populations) currently listed as Critically Endangered, Endangered and Vulnerable under the *EBPC Act* and applied our demographic framework.

First, we identified whether each taxon was considered at risk of fire-related threats in two comprehensive sources: the Australian Government Species Profile and Threats Database (hereafter ‘SPRAT’; https://www.environment.gov.au/cgi-bin/sprat/public/sprat.pl) and The Action Plan for Australian Mammals 2012 (hereafter ‘Action Plan’; Woinarski et al., 2014). These sources use several terms to describe fire-related threats including “inappropriate fire regimes”, “wildfire”, “change in fire regimes”, “modified fire regimes” and “habitat change due to altered fire regimes”.

Second, for each taxon considered at risk from a fire-related threat, we systematically reviewed two comprehensive sources and the primary literature. This included: (i) the SPRAT and related documentation (Conservation Advice, Listing Advice, Recovery Plan); (ii) taxon profiles in the Action Plan; (iii) primary literature cited in the SPRAT and Action Plan that underpinned the inclusion of fire as a threat and interactions with other processes (n = 164 papers); and (iv) additional primary literature (published 2010-2020) identified through systematic search in the Web of Science (n = 39). Recent primary literature was identified after harvesting key references from the SPRAT and Action Plan, and included searching the scientific name and common names of each taxon in Web of Science along with the terms *fire OR *burn* (see Supporting Information for details on all references analyzed in the systematic review).

Third, we used the information from all those sources to apply our demographic framework and identify the fire-driven mechanisms of decline, the associated fire-regime characteristics, and the processes that were considered to interact with fire. We considered variations in five fire-regime characteristics that could lead to mammalian population decline via ‘inappropriate’ fire regimes: 1 - *fire frequency* (high or low) – the number of fires in a given period; 2 - *fire intensity and severity* (high or low) – energy released from a fire (intensity) and its impact on plant biomass (severity); 3 - *fire patchiness* (uniform or patchy) – the configuration of post-fire landscapes including burns that are more uniform (i.e. coarse-scale patches of the same type) or patchy (i.e. fine-scale patches that are interspersed); 4 - *fire seasonality* (altered season) – consistently earlier or later peak flammability or longer periods of high flammability; and 5 - *fire size and amount* (large or small) – the size of fire events and the total amount of area burned by one or more fires. We then searched for evidence of links between the fire-regime characteristics and the seven mechanisms of decline described in our demographic framework.

We also recorded eight ecological processes that could interact with fire to affect mammals: 1 - climate and extreme weather (including linked changes in weather and climate such as pre- or post-fire drought and extreme fire weather); 2 - disease that directly influences animal populations; 3 - disease that influences habitat (i.e. vegetation dieback); 4 - grazing activity (including associated impacts of trampling and browsing by native or introduced herbivores); 5 - habitat loss and fragmentation; 6 - predation by introduced animals (e.g. cats, domestic dogs, foxes); 7 - predation by native animals (e.g. birds, dingoes, snakes); and 8 - weed invasion.

The quality of scientific evidence varies (Pullin & Knight, 2003) and preliminary analyses indicated that levels of evidence of fire-related threats differed among taxa and sources. Therefore, we established a method for classifying levels of evidence supporting reported fire-related threats and mechanisms of decline, and applied it systematically throughout the analysis (Table 1). We considered four types of empirical studies (manipulative experiments, longitudinal study, natural experiment and simulation modelling; following Driscoll et al., 2010a), as well as descriptive work, opinions of experts and anecdotal evidence, when assessing strength of evidence (see Figure S1 of Supporting Information for additional details). We acknowledge that deep knowledge of fire and mammals is held by local and Indigenous peoples across Australia. A limitation of the present review is that it is restricted to scientific literature and policy documents.

**TABLE 1.**
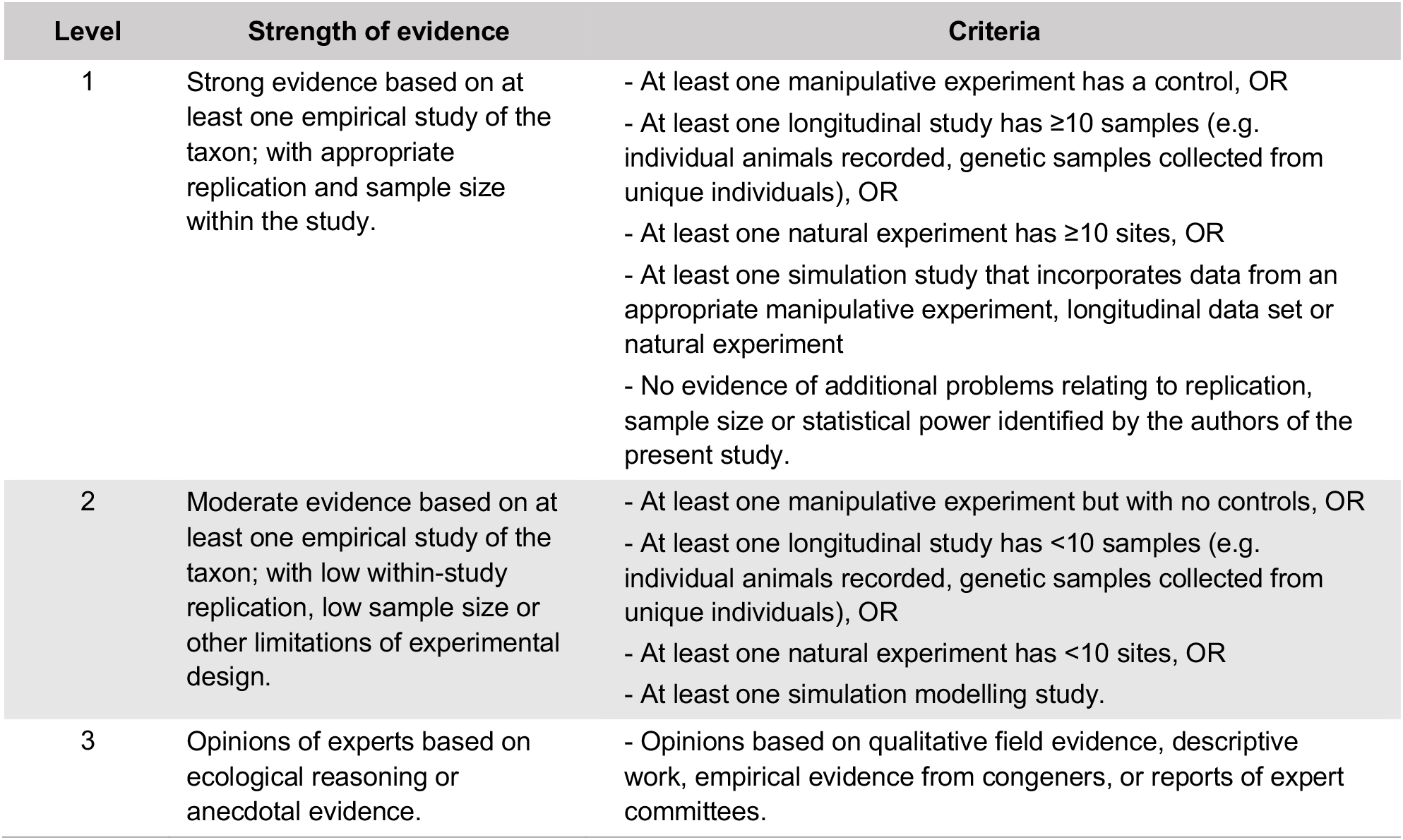
Levels of scientific evidence for classifying fire-driven mechanisms of decline, associated fire-regime characteristics and interacting ecological processes.

| Level | Strength of evidence | Criteria |
| --- | --- | --- |
| 1 | Strong evidence based on at least one empirical study of the taxon; with appropriate replication and sample size within the study. | - At least one manipulative experiment has a control, OR<br>- At least one longitudinal study has ≥10 samples (e.g. individual animals recorded, genetic samples collected from unique individuals), OR<br>- At least one natural experiment has ≥10 sites, OR<br>- At least one simulation study that incorporates data from an appropriate manipulative experiment, longitudinal data set or natural experiment<br>- No evidence of additional problems relating to replication, sample size or statistical power identified by the authors of the present study. |
| 2 | Moderate evidence based on at least one empirical study of the taxon; with low within-study replication, low sample size or other limitations of experimental design. | - At least one manipulative experiment but with no controls, OR<br>- At least one longitudinal study has <10 samples (e.g. individual animals recorded, genetic samples collected from unique individuals), OR<br>- At least one natural experiment has <10 sites, OR<br>- At least one simulation modelling study. |
| 3 | Opinions of experts based on ecological reasoning or anecdotal evidence. | - Opinions based on qualitative field evidence, descriptive work, empirical evidence from congeners, or reports of expert committees. |

To enable synthesis across taxa, we followed Van Dyck and Strahan’s (2008) groupings of species with similar taxonomy and ecology. We also recorded the vegetation types that each taxon inhabits, following vegetation classifications described by Keith (2017), according to information available in the primary literature, SPRAT and Action Plan (Tables S1 and S2).

## 3 RESULTS

### 3.1 Overview

Inappropriate fire regimes are listed as a threat to 88% (n = 87) of Australia’s terrestrial mammalian taxa listed as Critically Endangered, Endangered and Vulnerable under the *EPBC Act*. For 40% of those taxa (n = 35) the descriptions of fire-related threats were supported by empirical data (level of evidence 1 = 15%; n = 13; level of evidence 2 = 25%; n = 22) (Table S3). For 60% (n = 52) of the threatened terrestrial mammals considered to be at risk due to inappropriate fire regimes, the identification of fire-related threats was underpinned by opinions of experts based on ecological reasoning or anecdotal evidence (level of evidence 3).

A wide range of taxa have fire listed as a threat (Figure 2A). Moreover, there was evidence that fire threatens mammals inhabiting diverse vegetation types across Australia. For example, fire is a recorded threat for all threatened mammals that occur in hummock grasslands, savannas, and semi-arid eucalyptus woodlands (Figure 2B).

**FIGURE 2.**
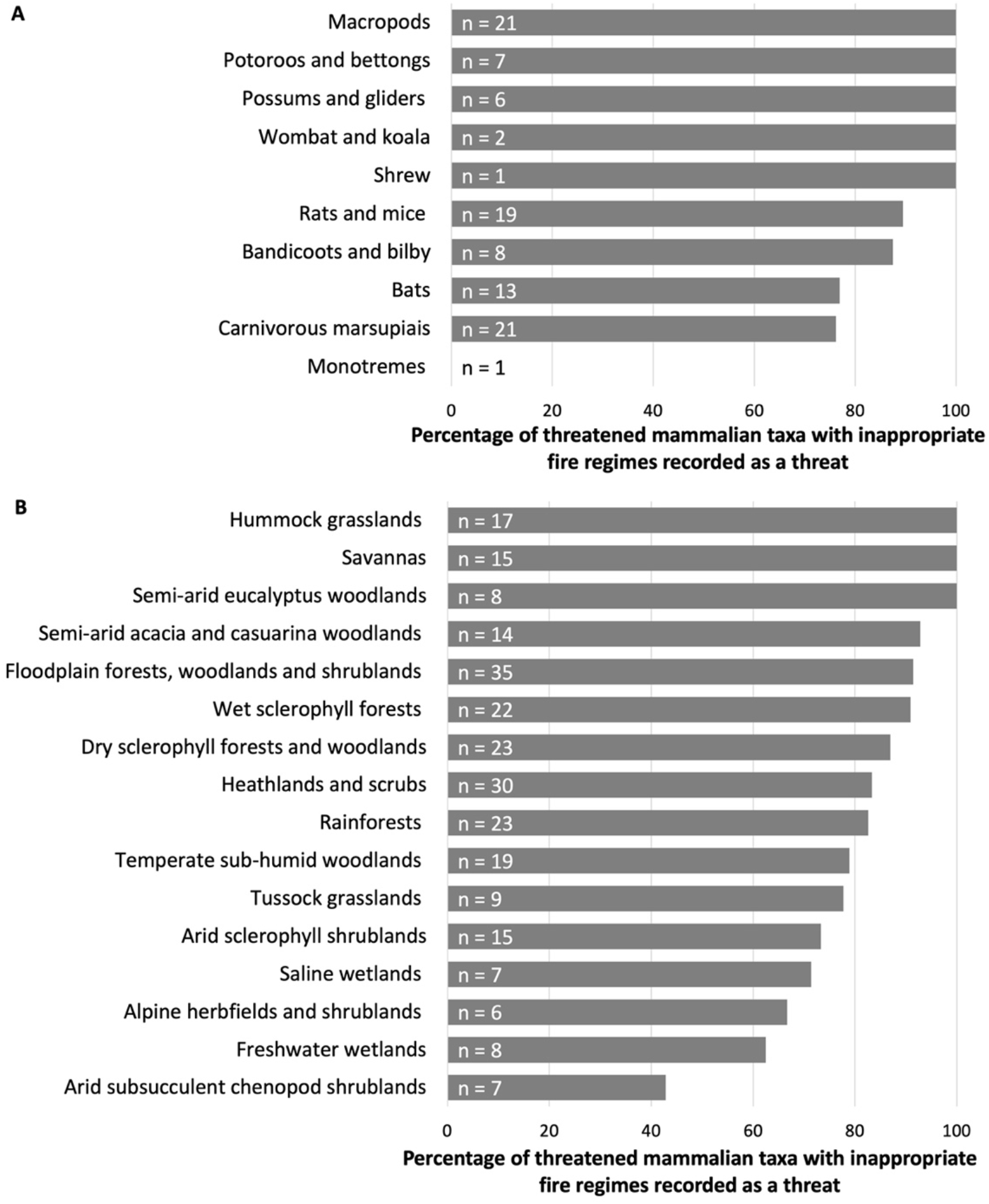
Percentage of Australian terrestrial mammalian taxa listed as Vulnerable, Endangered or Critically Endangered with inappropriate fire regimes recorded as a threat according to (A) taxonomic group and (B) vegetation type. n is the total number of threatened mammalian taxa listed as threatened in each taxonomic group (A) and vegetation type (B). Taxa may occur in more than one vegetation type.

### 3.2 Mechanisms of decline and fire-regime characteristics

The most frequently reported mechanism of fire-related decline was reduced survival shortly after fire due to reductions in food and shelter (M2; Figure 3A), which was identified for 94% (n = 82) of taxa threatened by inappropriate fire regimes. M2 was supported by a higher strength of evidence compared to other mechanisms: level of evidence 1 = 4% of cases identified as M2 (n = 3 taxa); level 2 = 28% (n = 23 taxa); and level 3 = 68% (n = 56 taxa). M2 was closely related to fire regimes with high fire frequency, high fire intensity and severity, and large fire size and amount (Figure 4).

**FIGURE 3.**
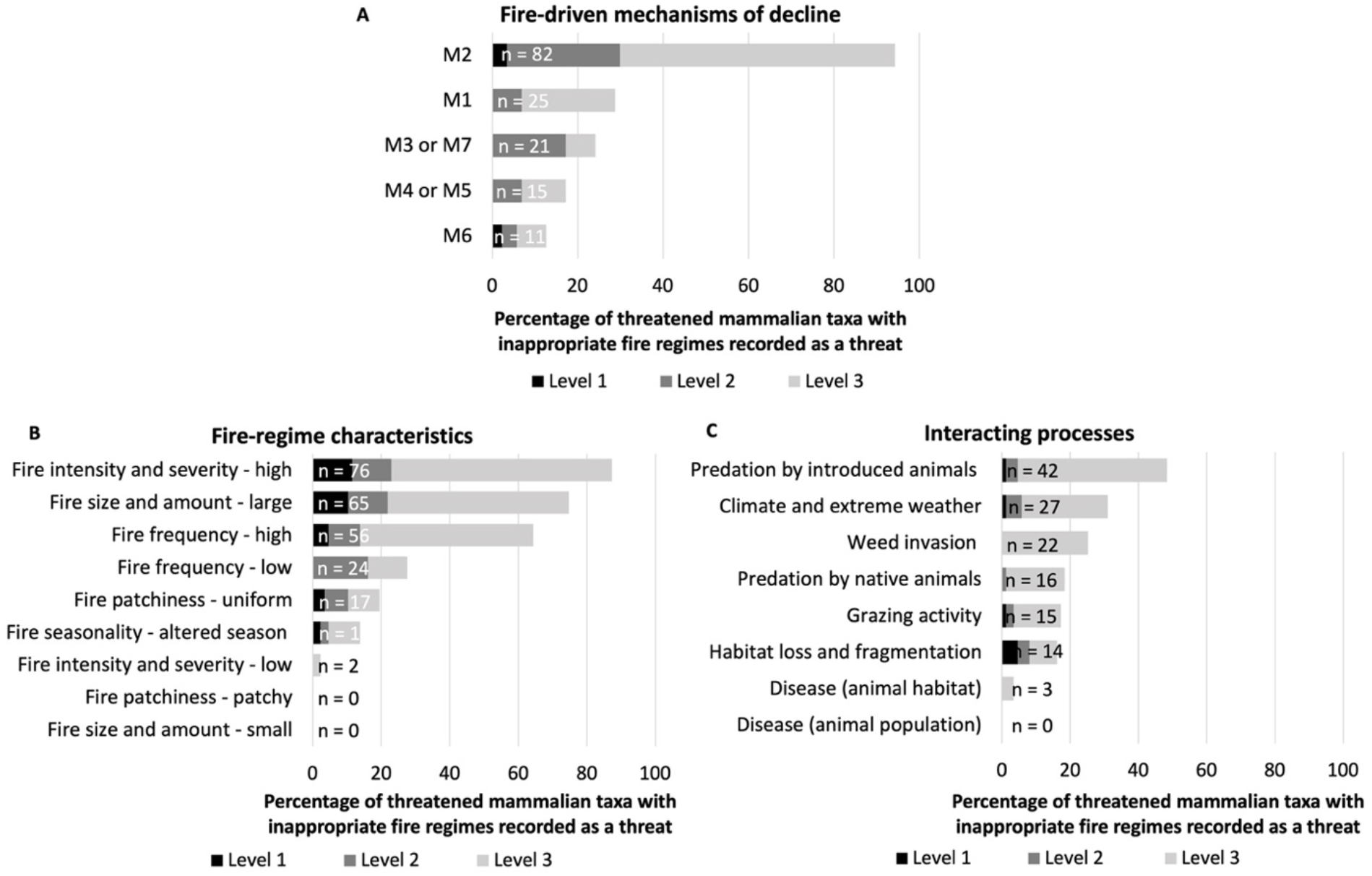
Percentage of Australian terrestrial mammalian taxa listed as Vulnerable, Endangered or Critically Endangered with inappropriate fire regimes recorded as a threat summarized by (A) Fire-driven mechanisms of decline; (B) Fire-regime characteristics; and (C) Interacting ecological processes. n is the total number of mammalian taxa threatened by fire for which the mechanisms, fire-regime characteristics or interacting processes were identified through systematic review. A given taxon can be affected by more than one mechanism, fire-regime characteristic or interacting process. Levels of evidence are shown by shading: Level 1 = Strong empirical evidence, Level 2 = Moderate empirical evidence, Level 3 = expert opinion and ecological reasoning (see Table 1 for descriptions of levels).

**FIGURE 4.**
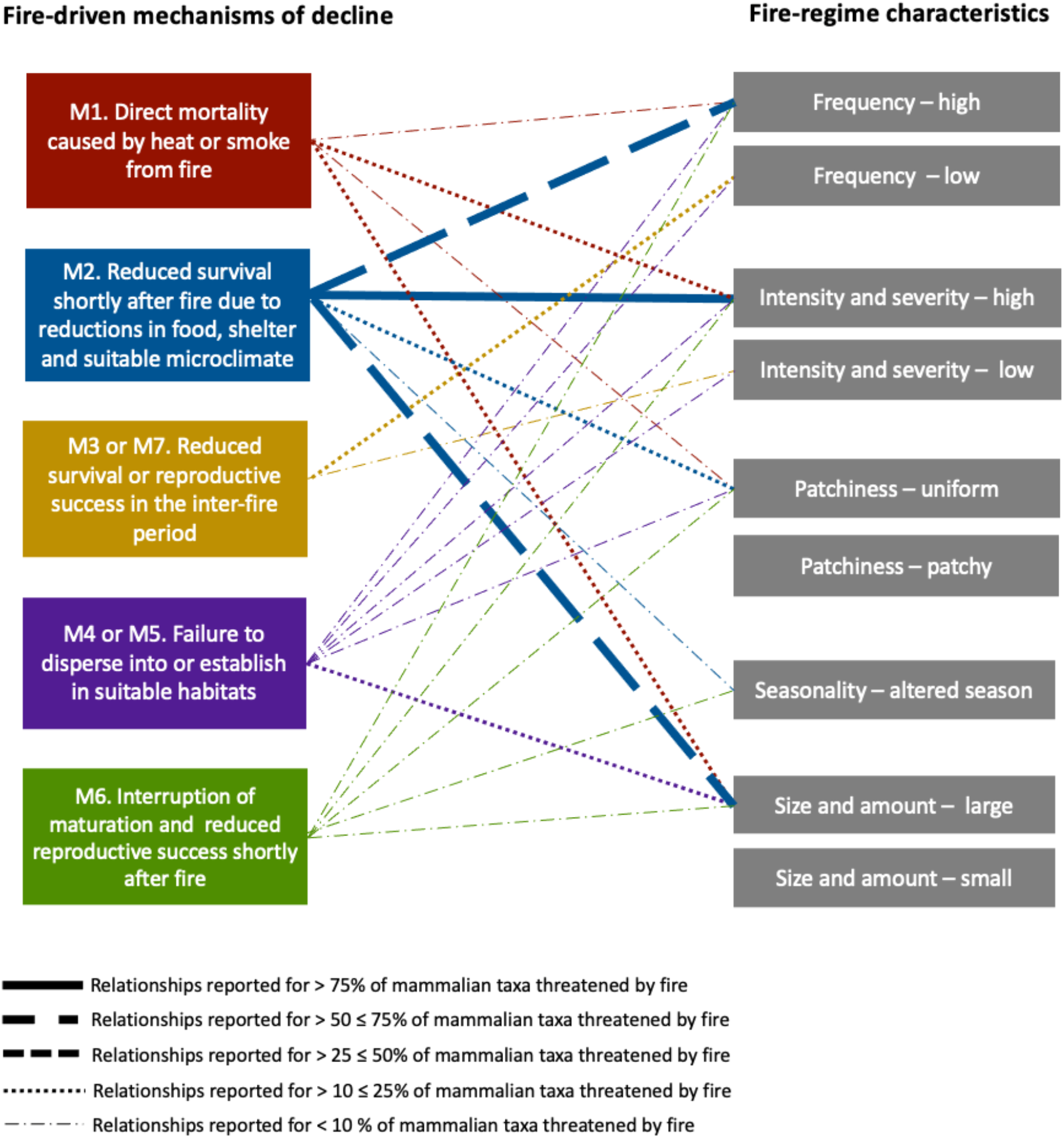
Relationships between fire-driven mechanisms of population decline (left) and fire-regime characteristics (right) identified for Australian terrestrial mammalian taxa listed as Vulnerable, Endangered or Critically Endangered with inappropriate fire regimes recorded as a threat. The percentage of taxa for which relationships between the mechanisms of decline and fire regime characteristics have been reported are indicated by different line types. Data are pooled across all levels of evidence.

Direct mortality caused by heat and smoke from fire (M1) was the second most frequent mechanism linked with mammal declines, affecting 29% (n = 25) of taxa threatened by fire (Figure 3A). The strength of evidence for M1 was low, with no cases based on level of evidence 1 (n = 0), 24% of cases (n = 6) based on level of evidence 2 and 76% (n = 19) on level 3. Direct mortality was primarily associated with high fire intensity and severity, and large fire size and amount (Figure 4).

For most taxa, reduced survival and reproduction following habitat change in the inter-fire period (M3 and M7, respectively) could not be separated based on available information. When combined, either M3 or M7 were inferred for 24% (n = 21) of the taxa threatened by fire (Figure 3A), with no instances of level of evidence 1 (n = 0). However, 71% (n = 15) of M3 and M7 were supported by level 2, and 29% (n = 6) by level 3. M3 and M7 were primarily linked to low fire frequency (Figure 4).

Reduced colonization rates in fire-prone ecosystems could not be pinpointed to being caused by either dispersal limitation (M4) or establishment limitation (M5) based on work to date. In total, M4 or M5 were inferred for 17% (n = 15) of taxa threatened by fire (Figure 3A), with no instances of level of evidence 1 (n = 0), 40% of the cases (n = 6) supported by level of evidence 2 and 60% (n = 9) by level of evidence 3. M4 and M5 were chiefly associated with large fire size (Figure 4).

Finally, interruption of maturation and reduced reproductive success shortly after fire (M6) was reported for only 13% of taxa (n = 11). 18% of the cases were classified as strong evidence (level 1; n= 2), 27% as moderate evidence (level 2; n = 3) and 55% based on expert opinion (level 3; n = 6). This mechanism was not clearly associated with a single fire-regime attribute (Figure 4).

Across all seven mechanisms of decline, inappropriate fire regimes were mostly characterized by high fire intensity and severity (87% of mammalian taxa threatened by fire; n = 76) (Figure 3B), large fire size and amount (75%; n = 65) and high fire frequency (64%; n = 56) (Figure 3B). Although measures associated with increased fire activity threaten more species, low fire frequency was a purported cause of inappropriate fire regimes for 28% (n = 24) of threatened mammals (Figure 3B). Patchy fires and small fires and were not reported as a threat to any taxa (Figure 3B) or associated with any of the seven main mechanisms of decline (Figure 4). For some taxa, the role of fire in causing population decline has been tested and, for the fire characteristics explored, considered negligible. That was the case for low fire intensity and severity (n = 6 taxa); patchy fires (n = 7), and small fire size and amount (n = 8). The southern brown bandicoot (*Isoodon obesulus obesulus*) is an example of species for which the effects of these fire regime characteristics on populations were considered negligible (Supporting Information).

### 3.3 Processes that interact with fire regimes

Seven of eight ecological processes were documented or predicted to interact with fire to exacerbate rates of mammalian decline. An association between fire and predation by introduced animals was the most frequently specified interaction, attributed to 48% (n = 42) of threatened mammalian taxa, followed by interactions with climate and extreme weather (25%; n = 27) and weed invasion (25%; n = 27) together. Levels of evidence were low for all interactions (Figure 3C).

## 4 DISCUSSION

Development and application of a demographic framework at a continental scale showed that inappropriate fire regimes for Australian mammals primarily comprise high-intensity and severe fire, large fire size and amount, and high-frequency fire. Each of these fire characteristics contribute to mammal declines primarily through reduced rates of survival. However, we also identified taxa for which inappropriate fire regimes include a frequency of fire that is too low. That is, some threatened mammalian populations are not getting enough of the ‘right’ kind of fire. Furthermore, systematic assessment of the levels of evidence underpinning fire-related declines indicated a lack of strong empirical evidence on relationships between demographic processes and fire-regime characteristics. The identification of these important knowledge gaps will help guide new work on animal survival, movement and dispersal in ecosystems that experience fire.

### 4.1 Defining inappropriate fire regimes

Our review of conservation assessments and primary literature showed that for most threatened Australian mammals, inappropriate fire regimes include high intensity and severity fires, large fire sizes and amount burned, and high-frequency fires. Intense and severe fires generate high levels of heat and smoke and increase the chances of animal mortality (Jolly et al., 2022). For example, declines of koala (*Phascolarctos cinereus*) populations have been documented following severe wildfires in forests of south-eastern Australia (Matthews et al., 2016; Phillips et al., 2021), and are probably linked to direct mortality caused by fires and reduced survival shortly after fires. Direct mortality caused by fire has been inferred for a range of mammals, including the western ringtail possum (*Pseudocheirus occidentalis*), red-tailed phascogale (*Phascogale calura*) and numbat (*Myrmecobius fasciatus*); however robust studies that directly measure mortality of threatened mammals during high intensity wildfires are rare (see Figure 3A and Supporting Information). Studies of experimental planned burns show that mortality of mammals can be low when fires are patchy and low intensity (e.g. Flanagan-Moodie et al., 2018; Vernes, 2000).

High intensity and severity fires can make habitat unsuitable for a range of threatened mammals by depleting critical food and shelter resources. For example, the abundance of greater gliders (*Petaurus volans*) declined after intense wildfires removed hollow-bearing trees and incinerated foliage, leading to scarcity of food and cover (Chia et al., 2015; Lindenmayer et al., 2013). Larger wildfires typically result in greater area burnt at high intensity and severity (Collins et al., 2021) and can cause widespread reductions in habitat and its connectivity. For instance, a population of swamp antechinus (*Antechinus minimus maritimus*) was considered extinct after a large (>40,000 ha) and severe wildfire in coastal heathlands of south-eastern Australia, with no recolonization detected in the subsequent 15 years (Wilson et al., 2018). Frequent fire can also be harmful: regular high severity fires in savannas of northern Australia have contributed to declines in populations of northern quolls (*Dasyurus hallucatus*) (Andersen, 2021; Griffiths & Brook, 2015), particularly through impacts on reproduction (Griffiths & Brook, 2015). While these and other examples (Supporting Information) are useful in examining characteristics that make a fire regime inappropriate, our results indicate that the impacts of fire regimes on survival, colonization, and reproduction are not documented by empirical evidence for many taxa (Figure 3A; Supporting Information). Further empirical studies will be crucial to reducing this uncertainty.

Why does a high frequency of intense and large fires pose a threat to many taxa that have evolved in Australian landscapes subject to recurrent fire? We propose the answer lies in concurrent changes to both mammalian populations and fire regimes. Several threatening processes, on their own and in combination, have reduced the size of mammalian populations, including habitat loss and fragmentation, and predation by introduced species (Doherty et al., 2015; Fisher et al., 2014; Woinarski et al., 2014). Smaller populations of mammals, which are restricted to increasingly narrow geographic areas, are then more likely to be harmed by intense and large fires. Examples of threatened mammals with small population sizes that are threatened by large and intense fires include Leadbeater’s possum (*Gymnobelideus leadbeateri*), Gilbert’s potoroo (*Potorous gilbertii*) and heath mouse (*Pseudomys shortridgei*). At the same time, there is increasing evidence that fire regimes are changing. For example, mega-fires in 2019-2020 burnt more than seven million ha across eastern Australia (Bowman et al., 2021). These fires were unprecedented in terms of their size and amount of area burned at high severity (Collins et al., 2021) and impacted the habitat of numerous threatened animals (Ward et al., 2020). Even ecosystems that have not historically experienced severe fires are experiencing increased activity of this type of fire e.g. the Gondwanan Rainforests of eastern Australia that are home to threatened mammals such as Hastings River mouse (*Pseudomys oralis*) and brush-tailed rock-wallaby (*Petrogale penicillata*) (Godfree et al., 2021).

Interestingly, a low frequency of fire was also an important characteristic of inappropriate fire regimes, documented or predicted to negatively affect 24 threatened mammal taxa. For example, reduced fire frequency leads to an alteration in vegetation structure in part of the northern bettong’s (*Bettongia tropica*) geographic range in northern Australia. In the absence of fire for long periods, rainforest-pioneering species dominate the understory, and litter cover accumulates, resulting in a reduction of important food resources for this potoroid (Abell et al., 2006; Bateman & Johnson, 2011). This can lead to population declines due low adult survival (M3) and low recruitment rates (M7). While the scarcity of early-successional habitats or fire-induced resources because of low frequency (or exclusion) of fire has been demonstrated to adversely affect the abundance of several mammals, including some species of macropods (Hayward et al., 2007), rodents (Davies et al., 2018) and arboreal marsupials (Trouvé et al., 2019), there is scant demographic information available to identify mechanisms of decline.

### 4.2 The biogeography of fire-related declines

Mammals are threatened by fire in a range of Australian ecosystems, from the arid interior of the continent to the coast and oceanic islands. Our systematic review indicates that no single taxonomic group is clearly at higher risk of extinction through inappropriate fire regimes; rather, extant threatened mammals are at risk from inappropriate fire regimes across the board: more than 75% of threatened species in each taxonomic group with more than one species are considered threatened by fire (Figure 2A). This result is consistent with risk assessments presented in the Action Plan for Australian Mammals, one source of data in the present review, which indicate that a wide range of threatened mammals, from different taxonomic groups and ecosystems, are at high risk of extinction from inappropriate fire regimes (Woinarski et al., 2014).

The combination of frequent, high severity and large fires have been documented to cause declines of mammalian populations in diverse ecosystems of Australia, including tropical savannas (Lawes et al., 2015), arid grasslands (Letnic & Dickman, 2006) and temperate forests (Lindenmayer et al., 2012). While less common, negative effects of reduced fire activity were also demonstrated for species inhabiting a range of different ecosystems, such as woodlands in north-eastern Australia (Jackson et al., 2020) and hummock grasslands in central Australia (Southgate & Carthew, 2006).

Nevertheless, our analyses point to some differences in how fire regimes threaten mammal populations in different regions. For example, the negative influence of altered fire season on mammal populations was more frequently documented in Australian tropical savannas. In these ecosystems, late-dry-season fires tend to be more intense and severe than early-dry-season fires. The timing and intensity of late-dry-season fires can affect reproduction of species such as northern quolls, which have a synchronous annual breeding cycle (Begg et al., 1981; Griffiths & Brook, 2015). There have been few studies on changes in fire seasonality in temperate areas and how they shape mammalian populations, and we encourage further empirical research on this topic. Another way forward would be to combine expert opinion and mathematical modelling to quantify the probability that changes in fire season, and other characteristics of fire regimes on their own or in combination, will drive species to extinction (Hayward, 2009).

### 4.3 Interactions between fire and other processes

A range of processes were documented to interact with fire and intensify mammal declines. Predation by introduced animals – particularly by red fox (*Vulpes vulpes*) and feral cat (*Felis catus*) – was the most frequent interacting process cited in peer-reviewed papers and policy documents. Introduced predators could exacerbate mammalian declines if hunting activity and/or hunting success increases in recently burned areas, due to the loss of understory vegetation leaving native mammals more exposed (Hradsky, 2020). While there is growing empirical evidence that cats and foxes increase hunting activity in recently burned areas (Hradsky et al., 2017; Leahy et al., 2015; McGregor et al., 2015), our review indicated limited empirical evidence on combined impacts of fire and predation on threatened mammals.

Climate and extreme weather were also identified as important processes interacting with fire to contribute to mammalian declines. The combination of extreme drought and severe fire weather contributed to the occurrence of the 2019-2020 mega-fires (Abram et al., 2021), which burnt more than 70% of habitat of the long-footed potoroo (*Potorous longipes*) (Geary et al., 2021). Post-fire drought was another climate-fire interaction identified in our systematic review. For example, native rodents of the genus *Pseudomys* may reach high numbers after periods of high rainfall following fire; however, when fire is followed by drought, vegetation grows slower and resources become scarce, compromising population recovery (Crowther et al., 2018; Hale et al., 2016).

We also identified a range of other processes that interact with fire to exacerbate mammal declines (Figure 3C) including habitat loss and fragmentation (e.g. western ringtail possum, *Pseudocheirus occidentalis*; Wayne et al., 2006), grazing activity (e.g. brush-tailed rock-wallaby, *Petrogale penicillata*; Tuft et al., 2012) and weed invasion (e.g. warru, *Petrogale lateralis*; Read and Ward, 2011). However, in policy documents, interactions between threats were often described as ‘potential’, reflecting scarcity of empirical data. New empirical studies focused on interactions between fire and emerging threats should be a priority for future research.

### 4.4 Implications for fire management and conservation policy

A framework that links changes in populations to fire-regime characteristics will help develop more effective conservation actions and policies (Figure 5) in a variety of global contexts. First, identifying the characteristics that describe inappropriate fire regimes for different species, and understanding the demographic processes underpinning population declines, help orient species-specific actions. Second, a focus on mechanisms helps recognize interactions that cause population declines and hence threats that need to be managed alongside fire. Figure 5 highlights examples of potential conservation actions informed by a demographic approach.

**FIGURE 5.**
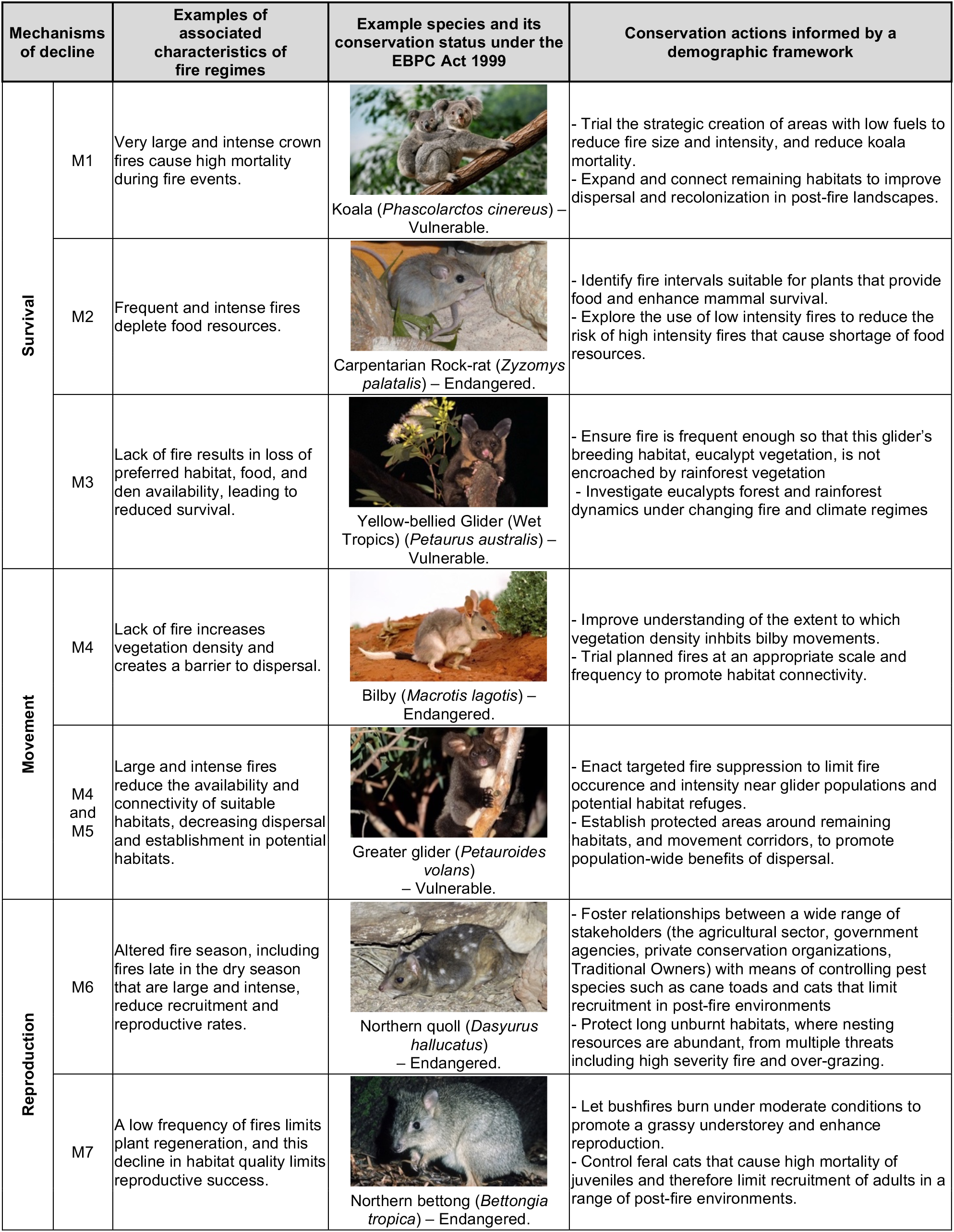
A demographic framework informs understanding of fire-driven population declines and conservation actions that could be taken to address them. The examples of conservation actions include some that have been implemented and others that have been proposed but not implemented. We recommend that actions be trialed and implemented through adaptive management that includes regular monitoring of mammal populations.

A range of emerging actions and strategies will be needed to manage fire for mammal conservation in Australia and globally. These include habitat restoration, Indigenous fire stewardship, planned burning, rapid recovery teams that assist wildlife after fire, reintroductions and targeted fire suppression (Bird et al., 2018; Geary et al., 2021; Martins et al., 2022; Roberts et al., 2022). Models that simulate management alternatives and different fire regimes offer opportunities to explore the effectiveness of potential strategies (Nitschke et al., 2020). Implementation through adaptive management and long-term monitoring is essential for determining which strategies will best promote populations of threatened mammals (Corey et al., 2020; Driscoll et al., 2010b).

## 5 CONCLUSION

There are exciting opportunities to apply the demographic framework we have developed to other taxa and regions. Recent work indicates that a range of animal taxa face threats related to changes in fire regimes, including amphibians, birds, dragonflies and damselflies, freshwater fishes and reptiles (Kelly et al., 2020). Changes in fire regimes are occurring worldwide, from arid, Mediterranean, temperate ecosystems to the tropics and tundra (Rogers et al., 2020). We anticipate that mechanistic approaches will help understand the causes and consequences of inappropriate fire regimes, and develop conservation policies and actions that address synergistic changes to the global environment.

## ACKNOWLEDGEMENTS

J.S was funded by the Holsworth Wildlife Research Endowment, Australian Wildlife Society University Research Grant, and Ecological Society of Australia Student Research Award. L.K was funded by a Centenary Research Fellowship at the University of Melbourne. This project was supported by the Australian Government’s National Environmental Science Program through the Threatened Species Recovery Hub. Credit photos: Koala (*Phascolarctos cinereus*) - Gerard Lacz/Shutterstock; Carpentarian Rock-rat (*Zyzomys palatalis*) - Damien Stanioch; Yellow-bellied Glider (*Petaurus australis*) - Josh Bowell; Bilby (*Macrotis lagotis*) - Michael Todd; Greater glider (*Petauroides volans*) - Josh Bowell; Northern quoll (*Dasyurus hallucatus*) - John Carnemolla/Shutterstock; Northern bettong (*Bettongia tropica*) - John Cancalosi/agefotostock.

