## Supporting Information for "Beyond inappropriate fire regimes: a synthesis of fire-driven declines of threatened mammals in Australia"

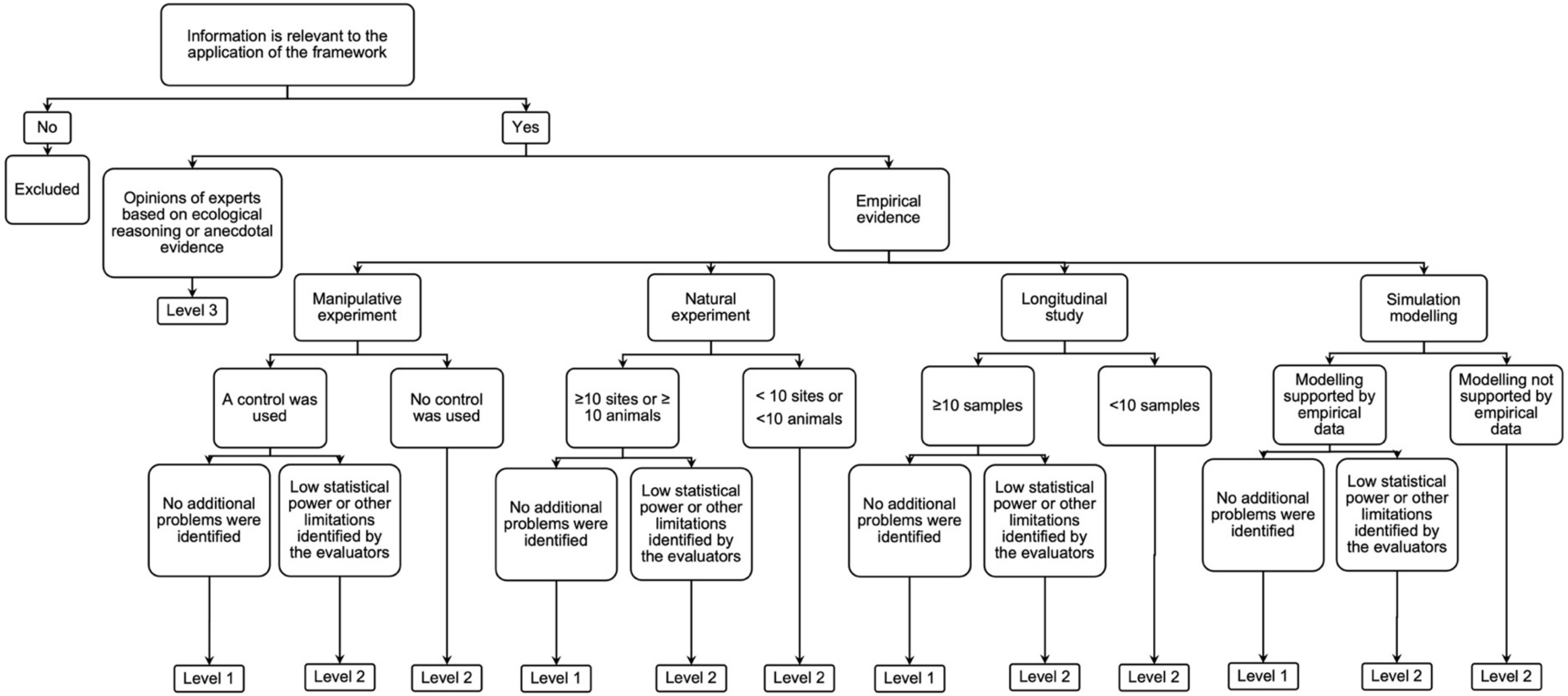


**FIGURE S1.** Decision tree for determining levels of evidence of fire-related population declines. Levels of evidence were used to classify fire-driven mechanisms of decline, fire-regime characteristics and interacting processes based on the information from primary literature and conservation assessments. We considered four types of empirical studies (manipulative experiment, longitudinal study, natural experiment and simulation modelling; following Driscoll et al., 2010[1]), as well as opinions of experts. Movement studies using radio-tracking or observations of behaviour could be grouped as either manipulative experiment, natural experiment or longitudinal study depending on the study design. Level 1 - Strong evidence based on at least one empirical study of the taxon, with appropriate replication and sample size within the study; 2 - Moderate evidence based on at least one empirical study of the taxon, with low within-study replication, low sample size or other limitations of experimental design; 3 - Opinions of experts based on qualitative field evidence, descriptive work, empirical evidence from congeners, or reports of expert committees.

[1] Driscoll, D. A., Lindenmayer, D. B., Bennett, A. F., Bode, M., Bradstock, R. A., Cary, G. J., ... & York, A. (2010). Fire management for biodiversity conservation: key research questions and our capacity to answer them. *Biological conservation*, *143*(9), 1928-1939.

**Table S1 –** List of the 99 terrestrial mammalian taxa listed as threatened under the *Environment Protection and Biodiversity Conservation Act 1999* (EPBC Act), retrieved from Australian Government Species Profile and Threats Database (SPRAT) on 29 of June 2020. Taxonomic groups follow Van Dyck and Strahan's (2008) grouping of species with similar taxonomy and ecology. Scientific names and common names follow SPRAT listing. The sequence of taxa follows the SPRAT listings, which present the taxa classified in each category of threatened species under the EPBC Act (respectively): CR = Critically Endangered; EN = Endangered; VU = Vulnerable. Taxa within each category are in alphabetical order.

| **Category of threatened species (EPBC Act)** | **Taxonomic group (Van Dyck and Strahan, 2008 )** | **Family** | **Scientific name** | **Common name** |
|---|---|---|---|---|
| CR | shrew | Soricidae | *Crocidura trichura* | Christmas Island Shrew |
| CR | possums and gliders | Petauridae | *Gymnobelideus leadbeateri* | Leadbeater's Possum |
| CR | wombat | Vombatidae | *Lasiorhinus krefftii* | Northern Hairy-nosed Wombat, |
| CR | bats | Miniopteridae | *Miniopterus orianae bassanii* | Southern Bent-wing Bat |
| CR | bats | Vespertilionidae | *Pipistrellus murrayi* | Christmas Island Pipistrelle |
| CR | macropods | Macropodidae | *Petrogale concinna concinna* | Nabarlek (Victoria River District) |
| CR | potoroos and bettongs | Potoroidae | *Potorous gilbertii* | Gilbert's Potoroo, Ngilkat |
| CR | possums and gliders | Pseudocheiridae | *Pseudocheirus occidentalis* | Western Ringtail Possum |
| CR | bats | Pteropodidae | *Pteropus natalis* | Christmas Island Flying-fox |
| CR | rats and mice | Muridae | *Zyzomys pedunculatus* | Central Rock-rat |
| EN | carnivorous marsupials | Dasyuridae | *Antechinus argentus* | Silver-headed Antechinus |
| EN | carnivorous marsupials | Dasyuridae | *Antechinus arktos* | Black-tailed Antechinus |
| EN | potoroos and bettongs | Potoroidae | *Bettongia penicillata* | Woylie |
| EN | potoroos and bettongs | Potoroidae | *Bettongia tropica* | Northern Bettong |
| EN | possums and gliders | Burramyidae | *Burramys parvus* | Mountain Pygmy-possum |
| EN | carnivorous marsupials | Dasyuridae | *Dasyurus hallucatus* | Northern Quoll |
| EN | carnivorous marsupials | Dasyuridae | *Dasyurus maculatus gracilis* (North QLD) | Spotted-tailed Quoll |
| EN | carnivorous marsupials | Dasyuridae | *Dasyurus maculatus maculatus* (mainland) | Spot-tailed Quoll |
| EN | carnivorous marsupials | Dasyuridae | *Dasyurus viverrinus* | Eastern Quoll |
| EN | bats | Hipposideridae | *Hipposideros inornatus* | Arnhem Leaf-nosed Bat |
| EN | bandicoots | Peramelidae | *Isoodon obesulus obesulus* | Southern Brown Bandicoot (eastern) |

| Category of threatened species (EPBC Act) | Taxonomic group (Van Dyck and Strahan, 2008 ) | Family | Scientific name | Common name |
|---|---|---|---|---|
| EN | macropods | Macropodidae | *Lagorchestes hirsutus* | Mala, Rufous Hare-Wallaby (Central Australia) |
| EN | rats and mice | Muridae | *Mesembriomys gouldii gouldii* | Black-footed Tree-rat |
| EN | carnivorous marsupials | Myrmecobiidae | *Myrmecobius fasciatus* | Numbat |
| EN | macropods | Macropodidae | *Onychogalea fraenata* | Bridled Nail-tail Wallaby |
| EN | carnivorous marsupials | Dasyuridae | *Parantechinus apicalis* | Dibbler |
| EN | bandicoots | Peramelidae | *Perameles bougainville bougainville* | Western Barred Bandicoot (Shark Bay) |
| EN | bandicoots | Peramelidae | *Perameles gunnii* | Eastern Barred Bandicoot (Mainland) Victorian subspecies |
| EN | possums and gliders | Petauridae | *Petaurus gracilis* | Mahogany Glider |
| EN | macropods | Macropodidae | *Petrogale coenensis* | Cape York Rock-wallaby |
| EN | macropods | Macropodidae | *Petrogale concinna canescens* | Nabarlek (Top End) |
| EN | macropods | Macropodidae | *Petrogale concinna monastria* | Nabarlek (Kimberley) |
| EN | macropods | Macropodidae | *Petrogale lateralis lateralis* | Black-flanked Rock-wallaby |
| EN | macropods | Macropodidae | *Petrogale persephone* | Proserpine Rock-wallaby |
| EN | potoroos and bettongs | Potoroidae | *Potorous longipes* | Long-footed Potoroo |
| EN | rats and mice | Muridae | *Pseudomys fumeus* | Smoky Mouse |
| EN | rats and mice | Muridae | *Pseudomys oralis* | Hastings River Mouse |
| EN | rats and mice | Muridae | *Pseudomys shortridgei* | Heath Mouse |
| EN | bats | Pteropodidae | *Pteropus conspicillatus* | Spectacled Flying-fox |
| EN | carnivorous marsupials | Dasyuridae | *Sarcophilus harrisii* | Tasmanian Devil |
| EN | carnivorous marsupials | Dasyuridae | *Sminthopsis griseoventer aitkeni* | Kangaroo Island Dunnart |
| EN | carnivorous marsupials | Dasyuridae | *Sminthopsis psammophila* | Sandhill Dunnart |
| EN | monotremes | Tachyglossidae | *Tachyglossus aculeatus multiaculeatus* | Kangaroo Island Echidna |
| EN | rats and mice | Muridae | *Zyzomys palatalis* | Carpentarian Rock-rat |
| VU | carnivorous marsupials | Dasyuridae | *Antechinus bellus* | Fawn Antechinus |
| VU | carnivorous marsupials | Dasyuridae | *Antechinus minimus maritimus* | Swamp Antechinus (mainland) |
| VU | potoroos and bettongs | Potoroidae | *Bettongia lesueur* | Burrowing Bettong |

| Category of threatened species (EPBC Act) | Taxonomic group (Van Dyck and Strahan, 2008 ) | Family | Scientific name | Common name |
|---|---|---|---|---|
| VU | potoroos and bettongs | Potoroidae | *Bettongia lesueur lesueur* | Burrowing Bettong (Shark Bay), Boodie |
| VU | bats | Vespertilionidae | *Chalinolobus dwyeri* | Large-eared Pied Bat, Large Pied Bat |
| VU | rats and mice | Muridae | *Conilurus penicillatus* | Brush-tailed Rabbit-rat |
| VU | carnivorous marsupials | Dasyuridae | *Dasyuroides byrnei* | Brushy-tailed marsupial rat |
| VU | carnivorous marsupials | Dasyuridae | *Dasyurus geoffroii* | Western Quoll, Chuditch |
| VU | carnivorous marsupials | Dasyuridae | *Dasyurus maculatus maculatus* | Spotted-tail Quoll (Tasmania) |
| VU | bats | Hipposideridae | *Hipposideros semoni* | Semon's Leaf-nosed Bat |
| VU | bandicoots | Peramelidae | *Isoodon auratus auratus* | Golden Bandicoot |
| VU | bandicoots | Peramelidae | *Isoodon auratus barrowensis* | Golden Bandicoot |
| VU | bandicoots | Peramelidae | *Isoodon obesulus nauticus* | Southern Brown Bandicoot |
| VU | macropods | Macropodidae | *Lagorchestes conspicillatus conspicillatus* | Spectacled Hare-wallaby (Barrow Isl) |
| VU | macropods | Macropodidae | *Lagorchestes hirsutus bernieri* | Rufous Hare-wallaby (Bernier Island) |
| VU | macropods | Macropodidae | *Lagorchestes hirsutus dorreae* | Rufous Hare-wallaby (Dorre Island) |
| VU | macropods | Macropodidae | *Lagostrophus fasciatus fasciatus* | Banded Hare-wallaby, Merrnine, Marnine, Munning |
| VU | rats and mice | Muridae | *Leporillus conditor* | Wopilkara, Greater Stick-nest Rat |
| VU | bats | Megadermatidae | *Macroderma gigas* | Ghost Bat |
| VU | bilby | Thylacomyidae | *Macrotis lagotis* | Greater Bilby |
| VU | rats and mice | Muridae | *Mastacomys fuscus mordicus* | Broad-toothed Rat (Mainland) |
| VU | rats and mice | Muridae | *Mesembriomys gouldii melvillensis* | Black-footed Tree-rat |
| VU | rats and mice | Muridae | *Mesembriomys gouldii rattoides* | Black-footed Tree-rat, Shaggy Rabbit-rat (N QLD) |
| VU | rats and mice | Muridae | *Notomys aquilo* | Northern Hopping-mouse, Woorrentinta |
| VU | rats and mice | Muridae | *Notomys fuscus* | Dusky Hopping-mouse, Wilkiniti |
| VU | bats | Vespertilionidae | *Nyctophilus corbeni* | Corben's Long-eared Bat, South-eastern Long-eared Bat |
| VU | macropods | Macropodidae | *Macropus robustus isabellinus* | Barrow Island Wallaroo, Barrow Island Euro |
| VU | bandicoots | Peramelidae | *Perameles gunnii gunnii* | Eastern Barred Bandicoot (Tasmania) |
| VU | possums and gliders | Pseudocheiridae | *Petauroides volans* | Greater Glider |

| Category of threatened species (EPBC Act) | Taxonomic group (Van Dyck and Strahan, 2008 ) | Family | Scientific name | Common name |
|---|---|---|---|---|
| VU | possums and gliders | Petauridae | *Petaurus australis* | Yellow-bellied Glider (Wet Tropics), Fluffy Glider |
| VU | macropods | Macropodidae | *Petrogale lateralis* | Warru, Black-footed Rock-wallaby (MacDonnell Ranges race) |
| VU | macropods | Macropodidae | *Petrogale lateralis* | Black-footed Rock-wallaby (West Kimberley race) |
| VU | macropods | Macropodidae | *Petrogale lateralis hacketti* | Recherche Rock-wallaby |
| VU | macropods | Macropodidae | *Petrogale penicillata* | Brush-tailed Rock-wallaby |
| VU | macropods | Macropodidae | *Petrogale sharmani* | Mount Claro Rock Wallaby, Sharman's Rock Wallaby |
| VU | macropods | Macropodidae | *Petrogale xanthopus celeris* | Yellow-footed Rock-wallaby (central-western Queensland) |
| VU | macropods | Macropodidae | *Petrogale xanthopus xanthopus* | Yellow-footed Rock-wallaby (SA and NSW) |
| VU | carnivorous marsupials | Dasyuridae | *Phascogale calura* | Red-tailed Phascogale, Red-tailed Wambenger, Kenngoor |
| VU | carnivorous marsupials | Dasyuridae | *Phascogale pirata* | Northern Brush-tailed Phascogale |
| VU | carnivorous marsupials | Dasyuridae | *Phascogale tapoatafa kimberleyensis* | Kimberley brush-tailed phascogale, Brush-tailed Phascogale (Kimberley) |
| VU | koala | Phascolarctidae | *Phascolarctos cinereus (combined populations of Qld, NSW and the ACT)* | Koala |
| VU | potoroos and bettongs | Potoroidae | *Potorous tridactylus tridactylus* | Long-nosed Potoroo (SE mainland) |
| VU | rats and mice | Muridae | *Pseudomys australis* | Plains Rat, Palyoora |
| VU | rats and mice | Muridae | *Pseudomys fieldi* | Shark Bay Mouse, Djoongari, Alice Springs Mouse |
| VU | rats and mice | Muridae | *Pseudomys novaehollandiae* | New Holland Mouse, Pookila |
| VU | rats and mice | Muridae | *Pseudomys pilligaensis* | Pilliga Mouse, Poolkoo |
| VU | bats | Pteropodidae | *Pteropus poliocephalus* | Grey-headed Flying-fox |
| VU | bats | Rhinolophidae | *Rhinolophus robertsi* | Large-eared Horseshoe Bat, Greater Large-eared Horseshoe Bat |
| VU | bats | Hipposideridae | *Rhinonicteris aurantia (Pilbara form)* | Pilbara Leaf-nosed Bat |
| VU | bats | Emballonuridae | *Saccolaimus saccolaimus nudicluniatus* | Bare-rumped Sheath-tailed Bat, Bare-rumped Sheathtail Bat |
| VU | macropods | Macropodidae | *Setonix brachyurus* | Quokka |
| VU | carnivorous marsupials | Dasyuridae | *Sminthopsis butleri* | Butler's Dunnart |
| VU | carnivorous marsupials | Dasyuridae | *Sminthopsis douglasi* | Julia Creek Dunnart |

| Category of threatened species (EPBC Act) | Taxonomic group (Van Dyck and Strahan, 2008 ) | Family | Scientific name | Common name |
|---|---|---|---|---|
| VU | rats and mice | Muridae | *Xeromys myoides* | Water Mouse, False Water Rat, Yirrkoo |
| VU | rats and mice | Muridae | *Zyzomys maini* | Arnhem Rock-rat, Arnhem Land Rock-rat, Kodjperr |

**Table S2 –** List of 99 terrestrial mammalian taxa listed as threatened under the *Environment Protection and Biodiversity Conservation Act 1999* (EPBC Act) and classification into the main vegetation types (following vegetation classifications described by Keith, 2017). Scientific names follow SPRAT listing (Australian Government Species Profile and Threats Database). Legend: 0 = absent; 1 = present. Vegetation types: **Rainfor** = Rainforests; **WetScler** = Wet sclerophyll forests; **DryScler** = sclerophyll forests and woodlands; **HeathScrub** = Heathlands and scrubs; **Savanna** = Savannas; **TempWood** = Temperate sub-humid woodlands; **TussGrass** = Tussock grasslands; **AlpShrub** = Alpine herbfields and shrublands; **FreshWet** = Freshwater wetlands; **FloodForest** = Floodplain forests, woodlands and shrublands; **SalineWet** = Saline wetlands; **SemiaridAcacia** = Semi-arid acacia and casuarina woodlands; **SemiaridEuc** = Semi-arid eucalyptus woodland; **AridChenop** = Arid subsucculent chenopod shrublands; **AridShrub** = Arid sclerophyll shrublands; **HummGrass** = Hummock grasslands; **Unknown** = Unknown/not classified.

| **Scientific name** | **Rainfor** | **WetScler** | **DryScler** | **HeathScrub** | **Savanna** | **TempWood** | **TussGrass** | **AlpShrub** | **FreshWet** | **FloodForest** | **SalineWet** | **SemiaridAcacia** | **SemiaridEuc** | **AridChenop** | **AridShrub** | **HummGrass** | **Unknown** |
|---|---|---|---|---|---|---|---|---|---|---|---|---|---|---|---|---|---|
| *Crocidura trichura* | 1 | 0 | 0 | 0 | 0 | 0 | 0 | 0 | 0 | 0 | 0 | 0 | 0 | 0 | 0 | 0 | 0 |
| *Gymnobelideus leadbeateri* | 0 | 0 | 0 | 0 | 0 | 1 | 0 | 0 | 0 | 0 | 0 | 0 | 0 | 0 | 0 | 0 | 0 |
| *Lasiorhinus krefftii* | 0 | 0 | 0 | 0 | 0 | 0 | 0 | 0 | 0 | 1 | 0 | 1 | 0 | 0 | 0 | 0 | 0 |
| *Miniopterus orianae bassanii* | 0 | 0 | 0 | 1 | 0 | 1 | 0 | 0 | 1 | 1 | 0 | 0 | 0 | 0 | 0 | 0 | 0 |
| *Pipistrellus murrayi* | 1 | 0 | 0 | 0 | 0 | 0 | 0 | 0 | 0 | 0 | 0 | 0 | 0 | 0 | 0 | 0 | 0 |
| *Petrogale concinna concinna* | 0 | 0 | 0 | 0 | 0 | 0 | 0 | 0 | 0 | 0 | 0 | 0 | 0 | 0 | 0 | 0 | 1 |
| *Potorous gilbertii* | 0 | 0 | 1 | 1 | 0 | 0 | 0 | 0 | 0 | 0 | 0 | 0 | 0 | 0 | 0 | 0 | 0 |
| *Pseudocheirus occidentalis* | 0 | 1 | 1 | 1 | 0 | 0 | 0 | 0 | 0 | 0 | 0 | 0 | 0 | 0 | 0 | 0 | 0 |
| *Pteropus natalis* | 1 | 0 | 0 | 0 | 0 | 0 | 0 | 0 | 0 | 0 | 0 | 0 | 0 | 0 | 0 | 0 | 0 |
| *Zyzomys pedunculatus* | 0 | 0 | 0 | 0 | 0 | 0 | 0 | 0 | 0 | 0 | 0 | 0 | 0 | 0 | 1 | 1 | 0 |
| *Antechinus argentus* | 0 | 1 | 1 | 0 | 0 | 0 | 0 | 0 | 0 | 0 | 0 | 0 | 0 | 0 | 0 | 0 | 0 |
| *Antechinus arktos* | 0 | 1 | 0 | 0 | 0 | 1 | 0 | 0 | 0 | 0 | 0 | 0 | 0 | 0 | 0 | 0 | 0 |
| *Bettongia penicillata* | 0 | 1 | 1 | 1 | 0 | 1 | 0 | 0 | 0 | 0 | 0 | 0 | 0 | 0 | 0 | 0 | 0 |
| *Bettongia tropica* | 0 | 1 | 0 | 0 | 0 | 0 | 0 | 0 | 0 | 0 | 0 | 0 | 0 | 0 | 0 | 0 | 0 |
| *Burramys parvus* | 0 | 0 | 0 | 0 | 0 | 0 | 0 | 1 | 0 | 0 | 0 | 0 | 0 | 0 | 0 | 0 | 0 |
| *Dasyurus hallucatus* | 1 | 1 | 1 | 1 | 1 | 0 | 0 | 0 | 0 | 1 | 0 | 1 | 0 | 0 | 0 | 0 | 0 |
| *Dasyurus maculatus gracilis* | 1 | 1 | 0 | 0 | 0 | 0 | 0 | 0 | 0 | 0 | 0 | 0 | 0 | 0 | 0 | 0 | 0 |
| *Dasyurus maculatus maculatus* | 1 | 1 | 1 | 1 | 0 | 1 | 0 | 0 | 0 | 1 | 0 | 0 | 0 | 0 | 0 | 0 | 0 |

| Scientific name | Rainfor | WetScler | DryScler | HeathScrub | Savanna | TempWood | TussGrass | AlpShrub | FreshWet | FloodForest | SalineWet | SemiaridAcacia | SemiaridEuc | AridChenop | AridShrub | HummGrass | Unknown |
|---|---|---|---|---|---|---|---|---|---|---|---|---|---|---|---|---|---|
| *Dasyurus viverrinus* | 0 | 0 | 1 | 1 | 0 | 1 | 1 | 1 | 0 | 0 | 0 | 0 | 0 | 0 | 0 | 0 | 0 |
| *Hipposideros inornatus* | 1 | 0 | 0 | 1 | 1 | 0 | 0 | 0 | 1 | 1 | 0 | 0 | 0 | 0 | 0 | 0 | 0 |
| *Isoodon obesulus obesulus* | 0 | 1 | 1 | 1 | 0 | 0 | 0 | 0 | 1 | 0 | 0 | 0 | 0 | 0 | 0 | 0 | 0 |
| *Lagorchestes hirsutus* | 0 | 0 | 0 | 0 | 0 | 0 | 0 | 0 | 0 | 0 | 0 | 0 | 0 | 0 | 0 | 1 | 0 |
| *Mesembriomys gouldii gouldii* | 0 | 0 | 0 | 0 | 0 | 0 | 0 | 0 | 0 | 1 | 0 | 0 | 0 | 0 | 0 | 0 | 0 |
| *Myrmecobius fasciatus* | 0 | 0 | 1 | 0 | 0 | 1 | 0 | 0 | 0 | 0 | 0 | 1 | 1 | 0 | 0 | 0 | 0 |
| *Onychogalea fraenata* | 0 | 0 | 0 | 0 | 0 | 0 | 0 | 0 | 0 | 0 | 0 | 1 | 1 | 0 | 0 | 0 | 0 |
| *Parantechinus apicalis* | 0 | 0 | 0 | 1 | 0 | 0 | 0 | 0 | 0 | 0 | 0 | 0 | 0 | 0 | 0 | 0 | 0 |
| *Perameles bougainville* | 0 | 0 | 0 | 0 | 0 | 0 | 0 | 0 | 0 | 0 | 0 | 0 | 0 | 1 | 1 | 0 | 0 |
| *Perameles gunnii* | 0 | 0 | 0 | 0 | 0 | 1 | 1 | 0 | 0 | 0 | 0 | 0 | 0 | 0 | 0 | 0 | 0 |
| *Petaurus gracilis* | 0 | 0 | 0 | 0 | 0 | 0 | 0 | 0 | 0 | 1 | 0 | 0 | 0 | 0 | 0 | 0 | 0 |
| *Petrogale coenensis* | 0 | 0 | 0 | 0 | 0 | 0 | 0 | 0 | 0 | 1 | 0 | 0 | 0 | 0 | 0 | 0 | 0 |
| *Petrogale concinna canescens* | 0 | 0 | 0 | 0 | 1 | 0 | 0 | 0 | 0 | 1 | 0 | 0 | 0 | 0 | 0 | 0 | 0 |
| *Petrogale concinna monastria* | 0 | 0 | 0 | 0 | 1 | 0 | 0 | 0 | 0 | 0 | 0 | 0 | 0 | 0 | 0 | 0 | 0 |
| *Petrogale lateralis lateralis* | 0 | 0 | 0 | 1 | 0 | 0 | 1 | 0 | 0 | 0 | 0 | 0 | 0 | 0 | 1 | 1 | 0 |
| *Petrogale persephone* | 1 | 0 | 0 | 0 | 0 | 0 | 0 | 0 | 0 | 1 | 0 | 0 | 0 | 0 | 0 | 0 | 0 |
| *Potorous longipes* | 1 | 1 | 1 | 1 | 0 | 1 | 0 | 0 | 0 | 1 | 0 | 0 | 0 | 0 | 0 | 0 | 0 |
| *Pseudomys fumeus* | 0 | 1 | 1 | 1 | 0 | 1 | 0 | 1 | 0 | 0 | 0 | 0 | 0 | 0 | 0 | 0 | 0 |
| *Pseudomys oralis* | 1 | 1 | 1 | 0 | 0 | 0 | 0 | 0 | 0 | 0 | 0 | 0 | 0 | 0 | 0 | 0 | 0 |
| *Pseudomys shortridgei* | 0 | 0 | 0 | 1 | 0 | 0 | 0 | 0 | 0 | 0 | 0 | 0 | 1 | 0 | 0 | 0 | 0 |
| *Pteropus conspicillatus* | 1 | 0 | 0 | 0 | 0 | 0 | 0 | 0 | 0 | 1 | 1 | 0 | 0 | 0 | 0 | 0 | 0 |
| *Sarcophilus harrisii* | 1 | 1 | 1 | 1 | 0 | 1 | 1 | 1 | 1 | 0 | 0 | 0 | 0 | 0 | 0 | 0 | 0 |
| *Sminthopsis griseoventer aitkeni* | 0 | 0 | 1 | 1 | 0 | 0 | 0 | 0 | 0 | 0 | 0 | 0 | 0 | 0 | 0 | 0 | 0 |
| *Sminthopsis psammophila* | 0 | 0 | 0 | 0 | 0 | 0 | 0 | 0 | 0 | 0 | 0 | 1 | 1 | 0 | 1 | 1 | 0 |
| *Tachyglossus aculeatus multiaculeatus* | 0 | 0 | 1 | 1 | 0 | 0 | 0 | 0 | 0 | 0 | 0 | 0 | 0 | 0 | 0 | 0 | 0 |
| *Zyzomys palatalis* | 1 | 0 | 0 | 0 | 1 | 0 | 0 | 0 | 0 | 1 | 0 | 0 | 0 | 0 | 0 | 0 | 0 |

| Scientific name | Rainfor | WetScler | DryScler | HeathScrub | Savanna | TempWood | TussGrass | AlpShrub | FreshWet | FloodForest | SalineWet | SemiaridAcacia | SemiaridEuc | AridChenop | AridShrub | HummGrass | Unknown |
|---|---|---|---|---|---|---|---|---|---|---|---|---|---|---|---|---|---|
| *Antechinus bellus* | 0 | 0 | 0 | 1 | 1 | 0 | 0 | 0 | 0 | 1 | 0 | 0 | 0 | 0 | 0 | 0 | 0 |
| *Antechinus minimus maritimus* | 0 | 1 | 0 | 1 | 0 | 0 | 1 | 0 | 0 | 1 | 0 | 0 | 0 | 0 | 0 | 0 | 0 |
| *Bettongia lesueur* | 0 | 0 | 0 | 0 | 0 | 0 | 0 | 0 | 0 | 0 | 0 | 1 | 1 | 1 | 1 | 1 | 0 |
| *Bettongia lesueur lesueur* | 0 | 0 | 0 | 0 | 0 | 0 | 0 | 0 | 0 | 0 | 0 | 1 | 1 | 1 | 1 | 1 | 0 |
| *Chalinolobus dwyeri* | 0 | 1 | 1 | 0 | 0 | 1 | 0 | 0 | 0 | 1 | 0 | 0 | 0 | 0 | 0 | 0 | 0 |
| *Conilurus penicillatus* | 0 | 0 | 0 | 0 | 1 | 0 | 0 | 0 | 0 | 1 | 0 | 0 | 0 | 0 | 0 | 0 | 0 |
| *Dasyuroides byrnei* | 0 | 0 | 0 | 0 | 0 | 0 | 0 | 0 | 1 | 0 | 0 | 0 | 0 | 1 | 1 | 0 | 0 |
| *Dasyurus geoffroii* | 0 | 1 | 1 | 1 | 0 | 0 | 0 | 0 | 0 | 1 | 0 | 0 | 0 | 0 | 0 | 0 | 0 |
| *Dasyurus maculatus maculatus* | 1 | 1 | 1 | 1 | 0 | 1 | 1 | 1 | 1 | 0 | 0 | 0 | 0 | 0 | 0 | 0 | 0 |
| *Hipposideros semoni* | 1 | 0 | 0 | 0 | 0 | 0 | 0 | 0 | 0 | 1 | 0 | 0 | 0 | 0 | 0 | 0 | 0 |
| *Isoodon auratus auratus* | 0 | 0 | 0 | 1 | 0 | 0 | 0 | 0 | 0 | 1 | 0 | 0 | 0 | 0 | 0 | 1 | 0 |
| *Isoodon auratus barrowensis* | 0 | 0 | 0 | 0 | 0 | 0 | 0 | 0 | 0 | 0 | 0 | 0 | 0 | 0 | 0 | 1 | 0 |
| *Isoodon obesulus nauticus* | 0 | 0 | 0 | 1 | 0 | 0 | 0 | 0 | 0 | 0 | 0 | 0 | 0 | 0 | 0 | 0 | 0 |
| *Lagorchestes conspicillatus conspicillatus* | 0 | 0 | 0 | 0 | 0 | 0 | 0 | 0 | 0 | 0 | 0 | 0 | 0 | 0 | 0 | 1 | 0 |
| *Lagorchestes hirsutus bernieri* | 0 | 0 | 0 | 0 | 0 | 0 | 0 | 0 | 0 | 0 | 0 | 0 | 0 | 0 | 0 | 1 | 0 |
| *Lagorchestes hirsutus dorreae* | 0 | 0 | 0 | 0 | 0 | 0 | 0 | 0 | 0 | 0 | 0 | 0 | 0 | 0 | 0 | 1 | 0 |
| *Lagostrophus fasciatus fasciatus* | 0 | 0 | 0 | 0 | 0 | 0 | 0 | 0 | 0 | 0 | 0 | 0 | 0 | 0 | 1 | 1 | 0 |
| *Leporillus conditor* | 0 | 0 | 0 | 1 | 0 | 0 | 0 | 0 | 0 | 0 | 0 | 0 | 0 | 1 | 0 | 0 | 0 |
| *Macroderma gigas* | 1 | 0 | 0 | 0 | 1 | 0 | 0 | 0 | 0 | 1 | 0 | 1 | 0 | 0 | 0 | 0 | 0 |
| *Macrotis lagotis* | 0 | 0 | 0 | 1 | 0 | 0 | 1 | 0 | 0 | 0 | 0 | 0 | 1 | 0 | 1 | 1 | 0 |
| *Mastacomys fuscus mordicus* | 0 | 0 | 0 | 0 | 0 | 0 | 0 | 1 | 0 | 1 | 0 | 0 | 0 | 0 | 0 | 0 | 0 |
| *Mesembriomys gouldii melvillensis* | 1 | 0 | 0 | 0 | 1 | 0 | 0 | 0 | 0 | 1 | 1 | 0 | 0 | 0 | 0 | 0 | 0 |
| *Mesembriomys gouldii rattoides* | 0 | 0 | 0 | 0 | 1 | 0 | 0 | 0 | 0 | 1 | 0 | 0 | 0 | 0 | 0 | 0 | 0 |
| *Notomys aquilo* | 0 | 0 | 0 | 0 | 1 | 0 | 0 | 0 | 0 | 1 | 0 | 0 | 0 | 0 | 0 | 0 | 0 |
| *Notomys fuscus* | 0 | 0 | 0 | 0 | 0 | 0 | 0 | 0 | 0 | 0 | 0 | 1 | 0 | 1 | 1 | 0 | 0 |
| *Nyctophilus corbeni* | 0 | 0 | 0 | 0 | 0 | 0 | 0 | 0 | 0 | 1 | 0 | 1 | 1 | 0 | 0 | 0 | 0 |

| Scientific name | Rainfor | WetScler | DryScler | HeathScrub | Savanna | TempWood | TussGrass | AlpShrub | FreshWet | FloodForest | SalineWet | SemiaridAcacia | SemiaridEuc | AridChenop | AridShrub | HummGrass | Unknown |
|---|---|---|---|---|---|---|---|---|---|---|---|---|---|---|---|---|---|
| *Macropus robustus isabellinus* | 0 | 0 | 0 | 0 | 0 | 0 | 0 | 0 | 0 | 0 | 1 | 0 | 0 | 0 | 0 | 1 | 0 |
| *Perameles gunnii gunnii* | 0 | 0 | 0 | 0 | 0 | 1 | 1 | 0 | 0 | 0 | 0 | 0 | 0 | 0 | 0 | 0 | 0 |
| *Petauroides volans* | 0 | 0 | 0 | 0 | 0 | 1 | 0 | 0 | 0 | 1 | 0 | 0 | 0 | 0 | 0 | 0 | 0 |
| *Petaurus australis* | 0 | 1 | 0 | 0 | 0 | 0 | 0 | 0 | 0 | 0 | 0 | 0 | 0 | 0 | 0 | 0 | 0 |
| *Petrogale lateralis* | 0 | 0 | 0 | 0 | 0 | 0 | 0 | 0 | 0 | 0 | 0 | 0 | 0 | 0 | 1 | 1 | 0 |
| *Petrogale lateralis* | 0 | 0 | 0 | 0 | 0 | 0 | 0 | 0 | 0 | 0 | 0 | 0 | 0 | 0 | 1 | 1 | 0 |
| *Petrogale lateralis hacketti* | 0 | 0 | 0 | 1 | 0 | 0 | 0 | 0 | 0 | 0 | 0 | 0 | 0 | 0 | 0 | 0 | 0 |
| *Petrogale penicillata* | 1 | 1 | 1 | 0 | 0 | 1 | 0 | 0 | 0 | 1 | 0 | 1 | 0 | 0 | 0 | 0 | 0 |
| *Petrogale sharmani* | 0 | 0 | 0 | 0 | 0 | 0 | 0 | 0 | 0 | 1 | 0 | 0 | 0 | 0 | 0 | 0 | 0 |
| *Petrogale xanthopus celeris* | 0 | 0 | 0 | 0 | 0 | 0 | 0 | 0 | 0 | 0 | 0 | 1 | 0 | 0 | 0 | 0 | 0 |
| *Petrogale xanthopus xanthopus* | 0 | 0 | 0 | 0 | 0 | 0 | 0 | 0 | 0 | 0 | 0 | 1 | 0 | 0 | 1 | 0 | 0 |
| *Phascogale calura* | 0 | 0 | 0 | 1 | 0 | 1 | 0 | 0 | 0 | 0 | 0 | 0 | 0 | 0 | 0 | 0 | 0 |
| *Phascogale pirata* | 0 | 0 | 0 | 0 | 1 | 0 | 0 | 0 | 0 | 1 | 0 | 0 | 0 | 0 | 0 | 0 | 0 |
| *Phascogale tapoatafa kimberleyensis* | 0 | 0 | 0 | 0 | 1 | 0 | 0 | 0 | 0 | 0 | 0 | 0 | 0 | 0 | 0 | 0 | 0 |
| *Phascolarctos cinereus* | 0 | 1 | 1 | 0 | 0 | 1 | 0 | 0 | 0 | 1 | 0 | 0 | 0 | 0 | 0 | 0 | 0 |
| *Potorous tridactylus tridactylus* | 0 | 1 | 1 | 1 | 0 | 0 | 0 | 0 | 0 | 0 | 0 | 0 | 0 | 0 | 0 | 0 | 0 |
| *Pseudomys australis* | 0 | 0 | 0 | 0 | 0 | 0 | 0 | 0 | 0 | 0 | 1 | 0 | 0 | 1 | 1 | 0 | 0 |
| *Pseudomys fieldi* | 0 | 0 | 0 | 0 | 0 | 0 | 0 | 0 | 0 | 0 | 1 | 0 | 0 | 0 | 0 | 1 | 0 |
| *Pseudomys novaehollandiae* | 1 | 0 | 1 | 1 | 0 | 1 | 0 | 0 | 0 | 0 | 0 | 0 | 0 | 0 | 0 | 0 | 0 |
| *Pseudomys pilligaensis* | 0 | 0 | 0 | 0 | 0 | 0 | 0 | 0 | 0 | 0 | 0 | 1 | 0 | 0 | 0 | 0 | 0 |
| *Pteropus poliocephalus* | 1 | 0 | 0 | 0 | 0 | 0 | 0 | 0 | 0 | 1 | 1 | 0 | 0 | 0 | 0 | 0 | 0 |
| *Rhinolophus robertsi* | 1 | 0 | 0 | 0 | 1 | 0 | 0 | 0 | 0 | 1 | 0 | 0 | 0 | 0 | 0 | 0 | 0 |
| *Rhinonicteris aurantia* | 0 | 0 | 0 | 0 | 0 | 0 | 0 | 0 | 0 | 1 | 0 | 0 | 0 | 0 | 1 | 0 | 0 |
| *Saccolaimus saccolaimus nudicluniatus* | 1 | 0 | 0 | 0 | 0 | 0 | 0 | 0 | 0 | 1 | 0 | 0 | 0 | 0 | 0 | 0 | 0 |
| *Setonix brachyurus* | 0 | 1 | 1 | 1 | 0 | 0 | 0 | 0 | 1 | 0 | 0 | 0 | 0 | 0 | 0 | 0 | 0 |
| *Sminthopsis butleri* | 0 | 0 | 0 | 0 | 1 | 0 | 0 | 0 | 0 | 0 | 0 | 0 | 0 | 0 | 0 | 0 | 0 |

| Scientific name | Rainfor | WetScler | DryScler | HeathScrub | Savanna | TempWood | TussGrass | AlpShrub | FreshWet | FloodForest | SalineWet | SemiaridAcacia | SemiaridEuc | AridChenop | AridShrub | HummGrass | Unknown |
|---|---|---|---|---|---|---|---|---|---|---|---|---|---|---|---|---|---|
| *Sminthopsis douglasi* | 0 | 0 | 0 | 0 | 0 | 0 | 1 | 0 | 0 | 0 | 0 | 0 | 0 | 0 | 0 | 0 | 0 |
| *Xeromys myoides* | 0 | 0 | 0 | 0 | 0 | 0 | 0 | 0 | 1 | 0 | 1 | 0 | 0 | 0 | 0 | 0 | 0 |
| *Zyzomys maini* | 1 | 0 | 0 | 0 | 0 | 0 | 0 | 0 | 0 | 0 | 0 | 0 | 0 | 0 | 0 | 0 | 0 |

**Table S3 –** List of 99 terrestrial mammalian taxa listed as threatened under the *Environment Protection and Biodiversity Conservation Act 1999* (EPBC Act), overall classification of the level of evidence for fire-related threats and bibliographic references analysed through systematic review (see Supporting Information – Database for detailed classification of fire-driven mechanisms of decline, fire-regime characteristics and interacting processes for each taxon and respective bibliographic references). Scientific names follow SPRAT listing (Australian Government Species Profile and Threats Database), for Critically Endangered, Endangered and Vulnerable taxa. Legend: **Level of evidence** (see Figure S1): **1** = Strong evidence based on at least one empirical study of the taxon; with appropriate replication and sample size within the study; **2** = Moderate evidence based on at least one empirical study of the taxon; with low within-study replication, low sample size or other limitations of experimental design; **3** = Opinions of experts based on ecological reasoning or anecdotal evidence. **NA** = not applicable (inappropriate fire regimes are not recorded as a threat). **References: 1 – References cited in the SPRAT and Action Plan for Australian Mammals 2012 (Woinarski et al., 2014)** = studies cited in the policy documents and that underpinned the description of fire-related threats and interacting processes; **2 – Additional references cited in 1** = relevant additional peer-reviewed papers resulted from unstructured search using the reference lists of studies cited in the SPRAT and Action Plan; **3 – Technical reports cited in the SPRAT and Action Plan** = technical reports, recovery plans and other references cited in the SPRAT and Action Plan; **4 – Web of Science 2010-2020** = relevant papers yielded from the systematic search of the peer-reviewed literature (published 2010 – 2020) in Web of Science and not cited in the SPRAT and the Action Plan. Searches were conducted in July 2020 using scientific names and common names of each terrestrial mammalian taxon listed under the EPBC Act along with the terms *fire OR *burn*. A complete list of all references follows this table. Note: taxa without references were assessed based on unpublished information from the SPRAT and Action Plan (e.g. expert opinion, unpublished reports).

| | | Bibliographic references | | | |
|---|---|---|---|---|---|
| **Scientific name** | **Level of evidence** | **1 - References cited in the SPRAT and Action Plan** | **2 - Additional references cited in 1** | **3 - Technical reports cited in the SPRAT and Action Plan** | **4 - Web of Science 2010 - 2020** |
| *Crocidura trichura* | 3 | | | | |
| *Gymnobelideus leadbeateri* | 1 | Lindenmayer et al. (2012); Lindenmayer et al. (2013a); Lindenmayer et al. (2017); Lindenmayer and Lacy (1995); Lindenmayer and Possingham (1995); Lindenmayer and Sato (2018); McBride et al. (2020); Nelson et al. (2019); Taylor et al. (2014); Taylor et al. (2017); Todd et al. (2016) | Banks et al. (2011); Banks et al. (2013); Lindenmayer et al. (2013b); Smith and Lindenmayer (1992) | Nelson et al. (2017) | Nitschke et al. (2020); Trouvé et al. (2019) |
| *Lasiorhinus krefftii* | 3 | | | Horsup (2004) | |
| *Miniopterus orianae bassanii* | NA | | | | |
| *Pipistrellus murrayi* | 3 | | | | |
| *Petrogale concinna concinna* | 3 | | | | |

| Scientific name | Level of evidence | Bibliographic references | | | |
|---|---|---|---|---|---|
| | | 1 - References cited in the SPRAT and Action Plan | 2 - Additional references cited in 1 | 3 - Technical reports cited in the SPRAT and Action Plan | 4 - Web of Science 2010 - 2020 |
| *Potorous gilbertii* | 3 | Nguyen and Friend (2005); Sinclair et al. (1996) | | | |
| *Pseudocheirus occidentalis* | 1 | Christensen and Abbot (1989); Inions (1985); Russell et al. (2003); Wayne et al. (2005); Wayne et al. (2006) | | Jones et al. (2004) | |
| *Pteropus natalis* | NA | | | | |
| *Zyzomys pedunculatus* | 2 | Edwards (2013); McDonald et al. (2013); McDonald et al. (2015); Nano et al. (2003) | | | Nano et al. (2019) |
| *Antechinus argentus* | 2 | Baker et al. (2013); Mason et al. (2017a) | | | Mason et al. (2017b) |
| *Antechinus arktos* | NA | | | | |
| *Bettongia penicillata* | 3 | Burrows and Christensen (2002); Marlow et al. (2015); Wayne et al. (2013) | Taylor (1991) | | Hing et al. (2017); Jones et al. (2018) |
| *Bettongia tropica* | 2 | Abell et al. (2006); Bateman and Johnson (2011); Harrington and Sanderson (1994); Vernes (2000); Vernes et al. (2001); Vernes and Haydon (2001); Vernes and Pope (2002) | Taylor (1991) | | Whitehead et al. (2018) |
| *Burramys parvus* | 1 | Broome et al. (2001); Mitrovski et al.(2008); Williams et al. (2008); Williams et al. (2012) | Qian et al. (2011) | | Gibson et al. (2018) |
| *Dasyurus hallucatus* | 1 | Begg et al. (1981); Friend and Taylor (1985); Oakwood (2000); Radford (2012); Woinarski et al. (2004a); Woinarski et al. (2004b); Woinarski et al. (2010) | Kitchener (1978) | | Griffiths and Brook (2015); Griffiths et al. (2015); Ibbet et al. (2018); Jolly et al. (2018); Radford and Andersen (2012) |
| *Dasyurus maculatus gracilis* | NA | | | | |
| *Dasyurus maculatus maculatus* | 1 | Dawson (2005); Dawson (2007); Glen and Dickman (2005) | Belcher (2007) | | |
| *Dasyurus viverrinus* | NA | | | | |
| *Hipposideros inornatus* | 3 | *Hipposideros inornatus* | | | |

| Scientific name | Level of evidence | Bibliographic references | | | |
|---|---|---|---|---|---|
| | | 1 - References cited in the SPRAT and Action Plan | 2 - Additional references cited in 1 | 3 - Technical reports cited in the SPRAT and Action Plan | 4 - Web of Science 2010 - 2020 |
| *Isoodon obesulus obesulus* | 2 | Arthur et al. (2012); Claridge and Barry (2000); Coates et al. (2008); Coates and Wright (2003) | Catling et al. (2001); Chia et al. (2016); Hale et al. (2016); Long (2009); McGregor et al. (2013); Paull (1995); Swan et al. (2015); Thompson et al. (1989) | New South Wales Department of Environment and Conservation (2006) | Hope et al. (2012); Ramalho et al. (2018) |
| *Lagorchestes hirsutus* | 2 | Bolton and Latz (1978); Lundie-Jenkins (1993) | | | |
| *Mesembriomys gouldii gouldii* | 2 | Friend et al. (1985); Friend (1987); Friend and Taylor (1985); Hohnen et al. (2015); Woinarski et al. (2004b) | | | Davies et al (2018a); Davies et al. (2018b) |
| *Myrmecobius fasciatus* | 3 | Christensen et al. (1984); Friend (1990) | Chirstensen and Abbott (1989) | | |
| *Onychogalea fraenata* | 3 | Fisher (2000) | | | |
| *Parantechinus apicalis* | 3 | | | Friend (2004) | |
| *Perameles bougainville bougainville* | 3 | Short et al. (1997); Short and Turner (1998) | | | |
| *Perameles gunnii* | 3 | Claridge and Barry (2000); Duft et al. (1991); Duft et al. (1994a); Duft et al. (1994b) | | | |
| *Petaurus gracilis* | 1 | Dettmann et al. (1995); Jackson et al. (1999); Jackson et al. (2000a); Jackson et al. (2000b); Jackson (2001); Jackson et al. (2011); Jackson and Claridge (1999); Tisdell et al. (2005) | | | |
| *Petrogale coenensis* | 3 | | | | |
| *Petrogale concinna canescens* | 3 | | | | |
| *Petrogale concinna monastria* | 3 | | | | |
| *Petrogale lateralis lateralis* | 3 | | | | |
| *Petrogale persephone* | 3 | | | | |

| Scientific name | Level of evidence | Bibliographic references | | | |
|---|---|---|---|---|---|
| | | 1 - References cited in the SPRAT and Action Plan | 2 - Additional references cited in 1 | 3 - Technical reports cited in the SPRAT and Action Plan | 4 - Web of Science 2010 - 2020 |
| *Potorous longipes* | 2 | Claridge and Barry (2000); Green et al. (1998) | Swan et al. (2015) | | |
| *Pseudomys fumeus* | 2 | Cockburn (1981); Ford et al. (2003) | | Nelson et al. (2009) | Burns et al. (2015) |
| *Pseudomys oralis* | 2 | Meek et al. (2003) | | | Law et al. (2016) |
| *Pseudomys shortridgei* | 2 | Di Stefano et al. (2011) | | | Di Stefano et al. (2014) |
| *Pteropus conspicillatus* | 3 | | | | |
| *Sarcophilus harrisii* | NA | | | | |
| *Sminthopsis griseoventer aitkeni* | 3 | | | Jones et al. (2010) in Gates et al. (2011) | |
| *Sminthopsis psammophila* | 2 | Miller et al. (2010) | | Churchill (2001) | Moseby et al. (2016) |
| *Tachyglossus aculeatus multiaculeatus* | NA | | | | |
| *Zyzomys palatalis* | 2 | Brook et al. (2002); Puckey et al. (2004); Trainor et al. (2000) | | | |
| *Antechinus bellus* | 2 | Begg et al. (1981); Friend (1985); Friend and Taylor (1985); Woinarski et al. (2004a); Woinarski et al. (2004b); Woinarski et al. (2010) | | | |
| *Antechinus minimus maritimus* | 1 | Magnusdottir et al. (2008); Wilson et al. (2001) | Taylor and Comfort (1993) | | Wilson et al. (2017) |
| *Bettongia lesueur* | 3 | Short et al. (1997); Short and Turner (1994) | | | |
| *Bettongia lesueur lesueur* | 3 | Short et al. (1997); Short and Turner (1994) | | | |
| *Chalinolobus dwyeri* | 3 | | | | |

| Scientific name | Level of evidence | Bibliographic references | | | |
|---|---|---|---|---|---|
| | | 1 - References cited in the SPRAT and Action Plan | 2 - Additional references cited in 1 | 3 - Technical reports cited in the SPRAT and Action Plan | 4 - Web of Science 2010 - 2020 |
| *Conilurus penicillatus* | 1 | Firth et al. (2006a); Firth et al. (2006b); Firth et al. (2010); Rossiter et al. (2003) | | | Davies et al (2018a); Davies et al. (2018b); Heiniger et al. (2020) |
| *Dasyuroides byrnei* | NA | | | | |
| *Dasyurus geoffroii* | 2 | Cardoso (2011); Morris et al. (2003) | | | |
| *Dasyurus maculatus maculatus* | 3 | Dawson (2005); Dawson (2007); Glen and Dickman (2005) | Belcher (2007) | | |
| *Hipposideros semoni* | 3 | | | | |
| *Isoodon auratus auratus* | 3 | Pardon et al. (2003) | | Palmer et al. (2003) | |
| *Isoodon auratus barrowensis* | 3 | Pardon et al. (2003) | | Palmer et al. (2003) | |
| *Isoodon obesulus nauticus* | NA | | | | |
| *Lagorchestes conspicillatus conspicillatus* | 3 | Ingleby (1991); Short and Turner (1991) | | | |
| *Lagorchestes hirsutus bernieri* | 3 | Bolton and Latz (1978); Lundie-Jenkins (1993); Richards et al. (2001); Richards and Short (1998); Short et al. (1997) | | | |
| *Lagorchestes hirsutus dorreae* | 3 | Bolton and Latz (1978); Lundie-Jenkins (1993); Richards et al. (2001); Richards and Short (1998); Short et al. (1997) | | | |
| *Lagostrophus fasciatus fasciatus* | 3 | Richards et al. (2001); Richards and Short (1998); Short et al. (1997) | | | |
| *Leporillus conditor* | 3 | | | | |
| *Macroderma gigas* | 3 | | | | |
| *Macrotis lagotis* | 2 | Moseby and O'Donnell (2003); Southgate et al. (2007); Southgate and Carthew (2006); Southgate and Carthew (2007) | | Bradley et al. (2015) | Cramer et al. (2016) |

| Scientific name | Level of evidence | Bibliographic references | | | |
|---|---|---|---|---|---|
| | | 1 - References cited in the SPRAT and Action Plan | 2 - Additional references cited in 1 | 3 - Technical reports cited in the SPRAT and Action Plan | 4 - Web of Science 2010 - 2020 |
| *Mastacomys fuscus mordicus* | 2 | Green and Osborne (2003); Green and Sanecki (2006); Hocking and Driessen (2000); Milner et al. (2015); O'Brien et al. (2008) | Taylor et al. (1985) | | |
| *Mesembriomys gouldii melvillensis* | 2 | Friend and Taylor (1985); Hohnen et al. (2015); Woirnarski et al. (2004b) | | | |
| *Mesembriomys gouldii rattoides* | 3 | Friend and Taylor (1985); Hohnen et al. (2015); Woirnarski et al. (2004b) | | | |
| *Notomys aquilo* | 3 | Woirnarski et al. (1999) | | | |
| *Notomys fuscus* | NA | Moseby et al. (2006) | | | |
| *Nyctophilus corbeni* | 3 | Parnaby et al. (2011) | | Parnaby et al. (2011) | |
| *Macropus robustus isabellinus* | 2 | Ealey (1967); Short and Turner (1991) | | | |
| *Perameles gunnii gunnii* | 3 | Claridge and Barry (2000) | | | |
| *Petauroides volans* | 1 | Andrew et al. 2014; Lindenmayer et al. (2000); Lindenmayer et al. (2011); Lindenmayer et al. (2013); Lunney (1987); Maloney (2007); McCarthy and Lindenmayer (1999a); McCarthy and Lindenmayer (1999b); Possingham et al. (1994); Taylor et al. (2007); Taylor and Goldingay (2005) | | | Berry et al. (2015); McLean et al. (2015); McLean et al. (2018); Smith and Smith (2018) |
| *Petaurus australis* | 3 | Lunney (1987); Eyre (2005) | Eyre et al. (2007) | | Eyre et al. (2010); Heise-Pavlov et al. (2017) |
| *Petrogale lateralis* | 3 | Muhic et al. (2012); Pearson et al. (1992); Read and Ward (2011); Ward et al. (2011) | West et al. (2017) | | |

| Scientific name | Level of evidence | Bibliographic references | | | |
|---|---|---|---|---|---|
| | | 1 - References cited in the SPRAT and Action Plan | 2 - Additional references cited in 1 | 3 - Technical reports cited in the SPRAT and Action Plan | 4 - Web of Science 2010 - 2020 |
| *Petrogale lateralis* | 3 | Muhic et al. (2012); Pearson et al. (1992); Read and Ward (2011); Ward et al. (2011) | West et al. (2017) | | |
| *Petrogale lateralis hacketti* | 3 | | | | |
| *Petrogale penicillata* | 1 | Tuft et al. (2012) | | | |
| *Petrogale sharmani* | 3 | | | | |
| *Petrogale xanthopus celeris* | 3 | Lapidge and Henshall (2001); Sharp and McCallum (2010) | | | |
| *Petrogale xanthopus xanthopus* | 3 | | | | |
| *Phascogale calura* | 2 | Kitchener (1981); Short and Hide (2012); Short et al. (2011) | | Friend and Wayne (2003) | |
| *Phascogale pirata* | 3 | Liedloff and Cook (2007); Wiliams et al. (1999) | | | |
| *Phascogale tapoatafa kimberleyensis* | 3 | Vigilante and Bowman (2004); van der Ree (2001) | | | |
| *Phascolarctos cinereus* | 1 | Gordon et al. (1988); Lunney et al. (2002); Lunney et al. (2004); Lunney et al. (2007); | | | Matthews et al. (2016) |
| *Potorous tridactylus tridactylus* | 1 | Norton et al. (2010a); Norton et al. (2010b) | Claridge et al., (1993) | | McHugh et al. (2019); McHugh et al. (2020) |
| *Pseudomys australis* | NA | | | | |
| *Pseudomys fieldi* | 3 | | | | |
| *Pseudomys novaehollandiae* | 2 | Braithwaite and Gullan (1978); Friend (1993); Fox (1982); Fox and Fox (1978); Fox and Mackay (1981); Keith and Calaby (1968); Seeback and Menkhorst (2000); Pye et al. (1991); Wilson (1991) | Kemper (1990); Crowther et al., (2018) | | Burns and Phillips (2020); Lock and Wilson (2017); Pedersen et al. (2014); Wilson et al. (2018) |

| Scientific name | Level of evidence | Bibliographic references | | | |
|---|---|---|---|---|---|
| | | 1 - References cited in the SPRAT and Action Plan | 2 - Additional references cited in 1 | 3 - Technical reports cited in the SPRAT and Action Plan | 4 - Web of Science 2010 - 2020 |
| *Pseudomys pilligaensis* | 2 | Tokushima et al. (2008); | Paull (2009); Paull et al. (2014); Tokushima and Jarman (2008); Tokushima and Jarman (2010); Tokushima and Jarman (2015) | | |
| *Pteropus poliocephalus* | 3 | | | | |
| *Rhinolophus robertsi* | 3 | | | | |
| *Rhinonicteris aurantia* | NA | | | | |
| *Saccolaimus saccolaimus nudicluniatus* | 3 | Murphy (2002) | | | |
| *Setonix brachyurus* | 1 | Burrows et al. (1995); Gibson et al.(2010); Hayward (2002); Hayward et al. (2004); Hayward et al. (2005a); Hayward et al. (2005b); Hayward (2007); Rippey and Hobbs (2003); De Tores et al. (2007) | Kitchener (1972) | | Bain et al. (2016); Dundas et al. (2018) |
| *Sminthopsis butleri* | 3 | | | | |
| *Sminthopsis douglasi* | 3 | | | | |
| *Xeromys myoides* | 3 | | | | |
| *Zyzomys maini* | 2 | Begg et al. (1981); Begg (1981); Kerle and Burgman (1984) | | | Ibbet et al. (2018); |

[2] The search 167 (conducted between 10 and 11 of July 2020) returned 173 papers distributed across 49 taxa. After excluding papers that were cited in the SPRAT and Action Plan (n = 27), and papers that were not relevant to  (n = 107), we identified 39 studies on 25 mammalian taxa that were not cited in the SPRAT and Action Plan and were included in the systematic review (see Table S3 in Supporting Information).

Griffiths, A. D., & Brook, B. W. (2015). Fire impacts recruitment more than survival of small-mammals in a tropical savanna. *Ecosphere*, 6(6), 1-22. doi: 10.1890/es14-00519.1

Griffiths, A. D., Garnett, S. T., & Brook, B. W. (2015). Fire frequency matters more than fire size: Testing the pyrodiversity–biodiversity paradigm for at-risk small mammals in an Australian tropical savanna. *Biological Conservation*, 186, 337-346. doi: 10.1016/j.biocon.2015.03.021

Heiniger, J., Davies, H. F., & Gillespie, G. R. (2020). Status of mammals on Groote Eylandt: Safe haven or slow burn?. *Austral Ecology*, 45(6), 759-772. doi: 10.1111/aec.12892

Heise-Pavlov, S. R., Chizinski, T., & Walker, N. E. (2017). Selection of sap feed trees by yellow-bellied gliders (*Petaurus australis*) in north-eastern Queensland, Australia–implications for site-specific habitat management. *Australian Mammalogy*, 40(1), 10-15. doi: 10.1071/am16035

Hing, S., Jones, K. L., Rafferty, C., Thompson, R. A., Narayan, E. J., & Godfrey, S. S. (2017). Wildlife in the line of fire: evaluating the stress physiology of a critically endangered Australian marsupial after bushfire. *Australian Journal of Zoology*, 64(6), 385-389. doi: 10.1071/zo16082

Hope, B. (2012). Short-term response of the long-nosed bandicoot, Perameles nasuta, and the southern brown bandicoot, *Isoodon obesulus obesulus*, to low-intensity prescribed fire in heathland vegetation. *Wildlife Research*, 39(8), 731-744. doi: 10.1071/wr12110

Ibbett, M., Woinarski, J. C. Z., & Oakwood, M. (2018). Declines in the mammal assemblage of a rugged sandstone environment in Kakadu National Park, Northern Territory, Australia. *Australian Mammalogy*, 40(2), 181-187. doi: 10.1071/am17011

Jolly, C. J., Kelly, E., Gillespie, G. R., Phillips, B., & Webb, J. K. (2018). Out of the frying pan: reintroduction of toad-smart northern quolls to southern Kakadu National Park. *Austral Ecology*, 43(2), 139-149. doi: 10.1111/aec.12551

Jones, K. L., Rafferty, C., Hing, S., Thompson, R. C. A., & Godfrey, S. S. (2018). Perturbations have minor impacts on parasite dynamics and body condition of an endangered marsupial. *Journal of Zoology*, 305(2), 124-132. doi: 10.1111/jzo.12541

Law, B., Brassil, T., & Gonsalves, L. (2016). Recent decline of an endangered, endemic rodent: does exclusion of disturbance play a role for Hastings River mouse (*Pseudomys oralis*)?. *Wildlife Research*, 43(6), 482-491. doi: 10.1071/wr16097

Lock, M., & Wilson, B. A. (2017). Influence of rainfall on population dynamics and survival of a threatened rodent (*Pseudomys novaehollandiae*) under a drying climate in coastal woodlands of south-eastern Australia. *Australian Journal of Zoology*, 65(1), 60-70. doi: 10.1071/zo16084

Mason, E. D., Firn, J., Hines, H. B., & Baker, A. M. (2017). Plant diversity and structure describe the presence of a new, threatened Australian marsupial within its highly restricted, post-fire habitat. *Plos one*, 12(8). doi: 10.1371/journal.pone.0182319

Matthews, A., Lunney, D., Gresser, S., & Maitz, W. (2016). Movement patterns of koalas in remnant forest after fire. *Australian Mammalogy*, 38(1), 91-104. doi: 10.1071/am14010

McHugh, D., Goldingay, R. L., Link, J., & Letnic, M. (2019). Habitat and introduced predators influence the occupancy of small threatened macropods in subtropical Australia. *Ecology and evolution*, *9*(11), 6300-6317. https://doi.org/10.1002/ece3.5203

McHugh, D., Goldingay, R. L., Parkyn, J., Goodwin, A., & Letnic, M. (2020). Short-term response of threatened small macropods and their predators to prescribed burns in subtropical Australia. *Ecological Management & Restoration*, 21(2), 97-107. doi: 10.1111/emr.12407

McLean, C. M., Bradstock, R., Price, O., & Kavanagh, R. P. (2015). Tree hollows and forest stand structure in Australian warm temperate Eucalyptus forests are adversely affected by logging more than wildfire. *Forest Ecology and Management*, 341, 37-44. doi: 10.1016/j.foreco.2014.12.023

McLean, C. M., Kavanagh, R. P., Penman, T., & Bradstock, R. (2018). The threatened status of the hollow dependent arboreal marsupial, the greater glider (*Petauroides volans*), can be explained by impacts from wildfire and selective logging. *Forest Ecology and Management*, 415, 19-25. doi: 10.1016/j.foreco.2018.01.048

Moseby, K., Read, J., McLean, A., Ward, M., & Rogers, D. J. (2016). How high is your hummock? The importance of *Triodia* height as a habitat predictor for an endangered marsupial in a fire-prone environment. *Austral Ecology*, 41(4), 376-389. doi: 10.1111/aec.12323

Nano, C. E., Randall, D. J., Stewart, A. J., Pavey, C. R., & McDonald, P. J. (2019). Spatio-temporal gradients in food supply help explain the short-term colonisation dynamics of the critically endangered central rock-rat (*Zyzomys pedunculatus*). *Austral Ecology*, 44(5), 838-849. doi: 10.1111/aec.12753

Nitschke, C. R., Trouvé, R., Lumsden, L. F., Bennett, L. T., Fedrigo, M., Robinson, A. P., & Baker, P. J. (2020). Spatial and temporal dynamics of habitat availability and stability for a critically endangered arboreal marsupial: implications for conservation planning in a fire-prone landscape. *Landscape Ecology*, 35, 1553-1570. doi: 10.1007/s10980-020-01036-2

Pedersen, S., Andreassen, H. P., Keith, D. A., Skarpe, C., Dickman, C. R., Gordon, I. J., ... & McArthur, C. (2014). Relationships between native small mammals and native and introduced large herbivores. *Austral Ecology*, 39(2), 236-243. doi: 10.1111/aec.12072

Radford, I. J., & Andersen, A. N. (2012). Effects of fire on grass-layer savanna macroinvertebrates as key food resources for insectivorous vertebrates in northern Australia. *Austral Ecology*, 37(6), 733-742. doi: 10.1111/j.1442-9993.2012.02413.x

Ramalho, C. E., Ottewell, K. M., Chambers, B. K., Yates, C. J., Wilson, B. A., Bencini, R., & Barrett, G. (2018). Demographic and genetic viability of a medium-sized ground-dwelling mammal in a fire prone, rapidly urbanizing landscape. *PloS one*, 13(2). doi: 10.1371/journal.pone.0191190

Smith, P., & Smith, J. (2018). Decline of the greater glider (*Petauroides volans*) in the lower Blue Mountains, New South Wales. *Australian Journal of Zoology*, 66(2), 103-114. doi: 10.1071/zo18021

Trouvé, R., Nitschke, C. R., Andrieux, L., Willersdorf, T., Robinson, A. P., & Baker, P. J. (2019). Competition drives the decline of a dominant midstorey tree species. Habitat implications for an endangered marsupial. *Forest Ecology and Management*, 447, 26-34. doi: 10.1016/j.foreco.2019.05.055

Whitehead, T., Vernes, K., Goosem, M., & Abell, S. E. (2018). Invasive predators represent the greatest extinction threat to the endangered northern bettong (*Bettongia tropica*). *Wildlife Research*, 45(3), 208-219. doi: 10.1071/wr16103

Wilson, B. A., Zhuang-Griffin, L., & Garkaklis, M. J. (2017). Decline of the dasyurid marsupial *Antechinus minimus maritimus* in south-east Australia: implications for recovery and management under a drying climate. *Australian Journal of Zoology*, 65(4), 203-216. doi: 10.1071/zo17041

Wilson, B. A., Lock, M., & Garkaklis, M. J. (2018). Long-term fluctuations in distribution and populations of a threatened rodent (*Pseudomys novaehollandiae*) in coastal woodlands of the Otway Ranges, Victoria: a regional decline or extinction?. *Australian Mammalogy*, 40(2), 281-293. doi: 10.1071/AM17036
